# *DPYD* links glial lipids and myelination to sleep duration

**DOI:** 10.64898/2026.10.31.763190

**Authors:** Amber J. Zimmerman, Samantha Deal, Erika Almeraya Del Valle, Joshua Donnelly, François Kroll, Xiaofeng Guo, Trisha R. Tsundupalli, Zoe Y. Shetty, Fusun Doldur-Balli, Brendan T. Keenan, Struan F.A. Grant, Jason Rihel, Amita Sehgal, Allan I. Pack

## Abstract

The regulation of sleep is heritable and conserved across species^1,2^, but the governing genetic mechanisms remain poorly understood. Here, we investigate the role of *DPYD*—which encodes the pyrimidine catabolic enzyme, dihydropyrimidine dehydrogenase—in sleep. Using a cross-species approach, we show that *DPYD* and its orthologs act specifically in glia to promote sleep through the regulation of brain lipids. In mouse and zebrafish, we find that *DPYD* orthologs are enriched in oligodendrocyte lineage cells, and their loss reduces myelination and sleep independent of *DPYD*’s established role in uracil catabolism to *β*-alanine. Genetic and pharmacological manipulations of myelin in zebrafish show that altering myelination is sufficient to perturb sleep duration. In *Drosophila*, which lack myelin, we find that the *DPYD* ortholog acts in ensheathing glia to promote sleep through formation of glial lipid droplets, a proposed evolutionary precursor to myelin. Furthermore, a human GWAS-identified sleep variant^2^ resides in a conserved cis-regulatory element that selectively contacts the promoter of *DPYD* within oligodendrocytes, connecting the human genetic association to oligodendrocyte-specific control of *DPYD*. Together, these results identify a deeply conserved role for glial lipids in sleep regulation and establish *DPYD* as a mechanistic link between myelin biology and sleep.

## Main

Sleep is a tightly controlled behavior that is conserved across species, yet the underlying mechanisms that govern sleep regulation are still not well understood^1^. Insufficient sleep is associated with myriad disorders, ranging from metabolic to neurocognitive^3–5^; thus, understanding the mechanisms that control sleep will inform therapeutic interventions for multiple disorders. Our recent work examined sleep and circadian behaviors in a large cohort of Diversity Outbred (DO) mice, a model that mimics human population genetic variation^6^. Using linkage mapping of quantitative sleep traits followed by phenotyping of genetic knockout mice, we identified a role for the *dihydropyrimidine dehydrogenase* gene (*DPYD*) in regulating sleep duration and consolidation^7,8^. Loss of *Dpyd* reduced sleep during the active phase in mice and fragmented sleep during the inactive phase^8^. However, the mechanism by which *DPYD* regulates sleep or, more broadly, cellular function in the central nervous system (CNS) is not known.

*DPYD* encodes the first and rate-limiting enzyme in pyrimidine catabolism, converting uracil and thymine to intermediate metabolites dihydrouracil and dihydrothymine. The reductive pyrimidine pathway provides cellular substrates, including *β*-alanine, that contribute to cellular metabolism^9,10^. DPYD deficiency is relatively common in humans due to numerous loss-of-function (LOF) genetic variants^11,12^, making DPYD a meaningful target for potential therapeutic intervention. Much of the work on this gene has been in the context of cancer research given that DPYD deficiency is associated with toxicity to the common chemotherapeutic 5-fluorouracil^13,14^ and that the chemotherapy adjuvant, gimeracil, inhibits DPYD function^15,16^. Outside of this, little is known of DPYD’s functional role.

To establish a cellular mechanism underlying the Dpyd-dependent sleep alterations, we designed a cross-species strategy in which different model systems addressed a distinct part of the causal chain. In mice, we localized *Dpyd* expression to oligodendrocyte lineage cells and showed that *Dpyd* loss reduces myelin thickness. Because myelin content and sleep are only associated in our mouse model, we used genetic and pharmacological manipulation of myelination in zebrafish to show that changing myelin content is sufficient to alter sleep and that restoring myelin can rescue sleep deficits caused by *DPYD* ortholog loss in this system. In *Drosophila*, which lack myelin entirely, we assessed the ancestral role of *DPYD* in glia, pointing to glial lipid droplet regulation as a candidate evolutionary precursor. Finally, we returned to human genetics to show that the sleep-duration variant at the *DPYD* locus is associated with oligodendrocyte-specific control of *DPYD* expression—implying the same regulatory mechanism identified across animal models. This cross-species design allowed us to address the fundamental role of *DPYD* in the control of sleep.

### Examining a role for Dpyd in the brain

Our previous work demonstrated that global loss of *Dpyd* in mice results in shorter sleep duration^8^, though the mechanism remains unresolved. Dpyd is the first and rate-limiting enzyme in pyrimidine catabolism to *β*-alanine (**Extended Data Fig. 1a**), and *β*-alanine has been proposed as an inhibitory neurotransmitter due to its structural similarity to GABA^17^; therefore, we first tested whether *Dpyd* loss altered brain *β*-alanine or related metabolites using liquid chromatography mass spectrometry (LC-MS). This experiment, however, revealed no significant differences in *β*-alanine levels in the brain or in any of the intermediate metabolites that are products of the other two pyrimidine catabolic enzymes, Dpys or Upb1 (**Extended Data Fig. 1b**). Uracil was significantly increased—as would be expected with loss of the catabolic enzyme—and the intermediate catabolite of Dpyd, dihydrothymine, was significantly decreased. Thus, Dpyd may be the only active enzyme in the pyrimidine catabolic pathway in the brain, which has been previously described^18^. We assayed other metabolites related to *β*-alanine including carnosine and histidine, along with neurotransmitters such as GABA and glutamate. None of these differed significantly between Dpyd knockouts and wild-type mice (**Extended Data Fig. 1c-d**), which is consistent with prior work showing there are other sources to these metabolites in the brain^9,17^. These results do not support the hypothesis that Dpyd affects sleep by altering *β*-alanine levels in the brain.

Other investigators have reported *Dpyd* as the only gene in the catabolic pathway expressed in the brain^18^; thus, we queried public single-cell RNA sequencing data from the *Tabula Muris* project^19^ to examine specific cell types expressing *Dpyd* that could be responsible for the sleep phenotype. As expected, all three genes encoding the catabolic enzymes *Dpyd*, *Dpys*, and *Upb1* are highly expressed in liver hepatocytes (**Extended Data Fig. 2a**). However, the cell type showing the highest *Dpyd* expression outside of hepatocytes is oligodendrocyte progenitor cells (OPCs) (**Extended Data Fig. 2a**). Neither *Dpys* nor *Upb1* are expressed in OPCs, suggesting these cells only utilize the metabolites generated by Dpyd: dihydrothymine and dihydrouracil. In the brain specifically, *Dpyd* showed little expression in other cell types (**Extended Data Fig. 2b**). These data suggest that Dpyd influences sleep through oligodendroglia in a manner that is independent of *β*-alanine synthesis.

In humans also, genetic variation at the *DPYD* locus is associated with sleep duration^20,21^; therefore, we reasoned that the cell type responsible for the sleep phenotype in both humans and mice would have higher expression of *DPYD* in both species. We queried brain-specific public single-cell RNA sequencing datasets from human^22^ and mouse^23^ to examine other brain cell types that may have been detected at lower levels in *Tabula Muris*. We observed that *DPYD* orthologs are expressed most abundantly in oligodendrocyte lineage cells in both human and mice (**Fig. 1a**). Given these are the myelinating cells of the central nervous system, we posited that loss of *Dpyd* could affect myelination.

**Figure 1.**
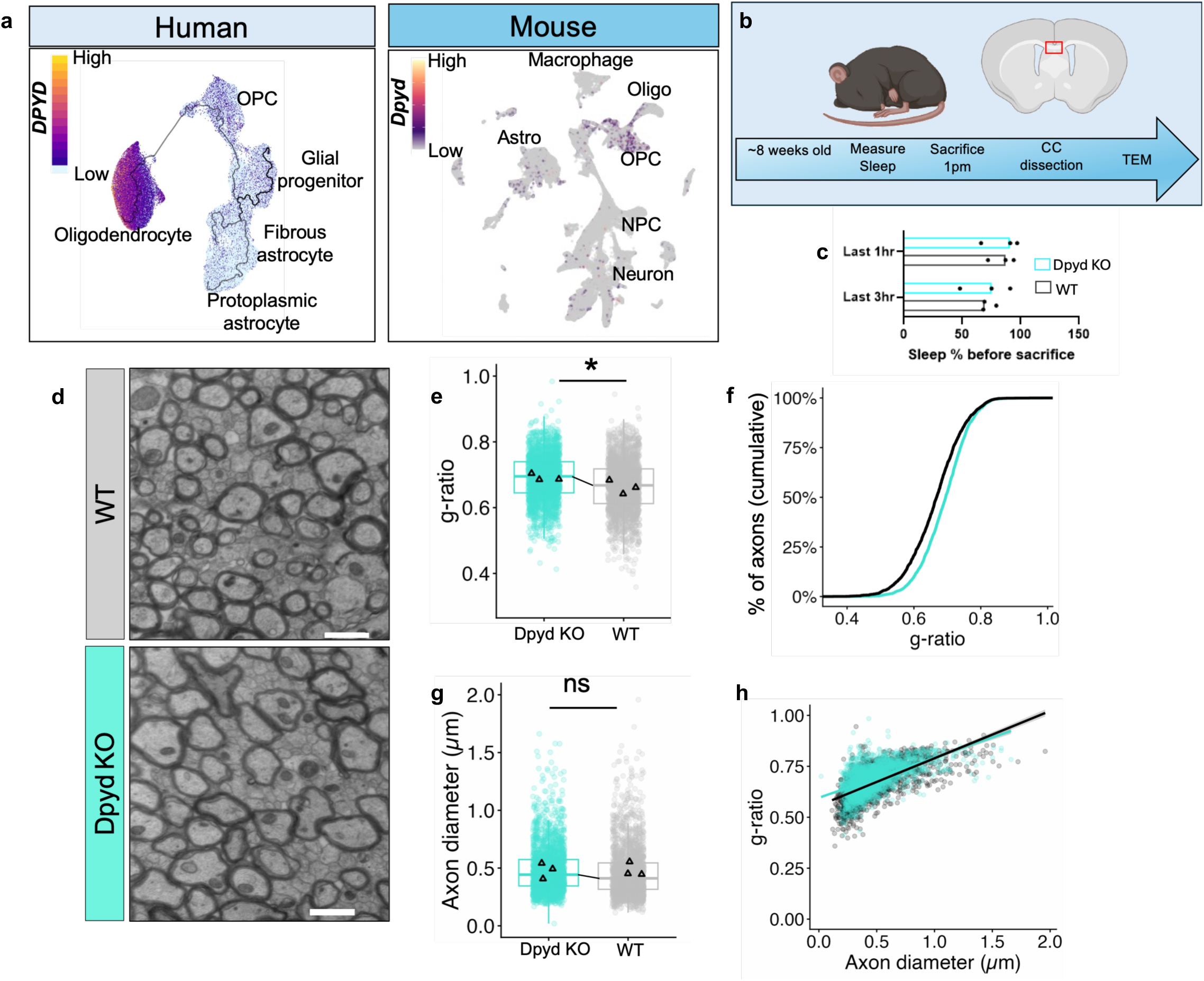
*Dpyd* is expressed in oligodendrocytes and affects myelination. **a.** UMAPs showing single-cell RNA expression of *DPYD* orthologs in human^22^ and mouse^23^ brain cells. Oligo = oligodendrocyte, OPC = oligodendrocyte progenitor cell, NPC = neural progenitor cell, Astro = astrocyte. **b.** Schematic showing timeline of mouse sleep analysis and brain corpus callosum (CC) dissection for transmission electron microscopy (TEM). **c.** Sleep percentage during the 1-hour and 3-hour block prior to sacrifice [n = 3 *Dpyd* KO and 3 wild-type (WT)]. **d.** Representative TEM images from *Dpyd* WT and *Dpyd* homozygous knockout (KO) mice showing cross section of myelinated axons in the corpus callosum. scale bar = 1µm. **e.** Genotype significantly influenced g-ratio (likelihood ratio test: χ²(1) = 4.79, *P* = 0.029), indicating thinner myelination in *Dpyd* KO mice. **f.** Cumulative frequency plot for g-ratio. **g.** No effect was observed for axon diameter (likelihood ratio test: χ²(1) = 0.004, *P* = 0.948). **h.** Scatter plot of g-ratio relative to axon diameter indicates a significant genotype X diameter interaction (t = 2.85, *P* = 0.004).

Oligodendrocyte precursors have been shown to proliferate during sleep and differentiate during wake^24^; therefore, we designed an experiment to delineate changes to myelin that were not driven by acute sleep/wake history or circadian differences. We assessed sleep using non-invasive infrared beam breaks based on a validated algorithm^25^ and sacrificed mice 6 hours into their sleep phase (Lights On) (**Fig. 1b**). Previous work indicated sleep percentage is highest at this time^26^ and total sleep duration only differed during the active phase in *Dpyd* knockouts, not during the first 6 hours of the sleep phase^8^. Indeed, we did not observe significant differences in the percent of time spent alseep in the 1-hour or 3-hour window prior to sacrifice between controls and *Dpyd* knockouts (**Fig. 1c**), allowing us to capture myelin changes impacted by genotype differences rather than acute sleep loss. Following sacrifice, we dissected the medial corpus callosum and measured myelin thickness using Transmission Electron Microscopy (TEM) (**Fig. 1d**). We calculated the g-ratio, where larger values indicate thinner myelin. Axons from *Dpyd* knockout mice displayed a significantly higher g-ratio relative to sibling wild-type controls (likelihood ratio test: χ²(1) = 4.79, *P* = 0.029) (**Fig. 1e-f**), indicative of thinner myelin. Axon diameter did not differ between groups (**Fig. 1g**), although the effect of genotype on the g-ratio was more prominent in smaller axons, as indicated by a significant interaction between genotype and axon diameter (t = 2.85, *P* = 0.004)

(**Fig. 1h**). These data implicate a role for *Dpyd* in oligodendrocyte myelination, suggesting a potential mechanism for the effect of *DPYD* on sleep duration.

### Zebrafish *dpyda/b* crispant phenocopies the mouse mutant

Our data from mice show that *Dpyd* loss results in shorter sleep and thinner myelin; however, the question remains whether the thinner myelin in *Dpyd* knockouts is secondary to the chronic short sleep or causative of shorter sleep. Several reports have determined sleep restriction reduces myelin across species^27–30^; though, whether the converse is true (i.e. myelin content has a direct effect on sleep duration) remains to be elucidated. Therefore, we sought to determine if directly altering myelination could influence sleep independent of direct loss of *DPYD*. Because genetic manipulation of myelin in mice is costly and likely to result in developmental lethality^31^, we turned to zebrafish, where genetic regulators of myelination are well established and do not result in grossly abnormal development^32,33^ that may confound sleep assessment. Zebrafish display diurnal sleep/wake rhythms similar to humans^34,35^ and have a rapid developmental timeline that facilitates genetic manipulation of myelin^36–42^, allowing for interrogation of behavior in multiple genetic mutants.

To first determine the validity of zebrafish as a model system for testing our hypotheses, we examined whether the *DPYD* ortholog is conserved in expression and function compared to mouse. In zebrafish, this gene is duplicated into *dpyda* and *dpydb*. Reanalysis of public single-cell RNA sequencing data from Daniocell^43^ revealed that *dpyda* is predominantly expressed in glia, while *dpydb* is expressed in the liver (**Fig. 2a**). While a metabolic role for DPYD in the liver has been described^16,44^ its expression has not been characterized in the zebrafish brain. To localize *dpyda/b* in the brain to a specific cell population, we examined glial subtypes from the Daniocell dataset, which showed *dpyda* is most abundantly expressed in the radial glia of the brain and spinal cord, while *dpydb* exhibits negligible expression in these cells (**Fig. 2b**). To visualize *dpyda* RNA expression in a whole animal, we performed *in situ* hybridization chain reaction (HCR) within whole-mount zebrafish larvae at 6 days post fertilization (dpf). We selected two marker genes from the single-cell data that labeled the glial population with high *dpyda* expression, *sox2* and *ptprz1b,* for *in vivo* co-labeling (**Fig. 2b-c**). Both are established radial glia markers^45^, with *ptprz1b* more specifically known to label oligodendrocyte precursor cells^46^. HCR was performed on *Tg*(*elavl3*:*H2B-GCaMP6s*) fish, which fluorescently label neuronal nuclei^47^. Consistent with the single cell data, *dpyda* was localized to ventricular domains (**Fig. 2c-f**), which are known locations of glial progenitors^48^. Specifically, *dpyda* was highly expressed in a region of the ventral telencephalon consistent with the progenitor population lining the telencephalic ventricle^49–51^. *dpyda* colocalized with *sox2* and *ptprz1b* within the ventricular domain and was largely absent from neuronal nuclei and other neuron-rich regions (**Fig. 2c-f**). Together, these data localized *dpyda* within the zebrafish brain to radial glia.

**Figure 2.**
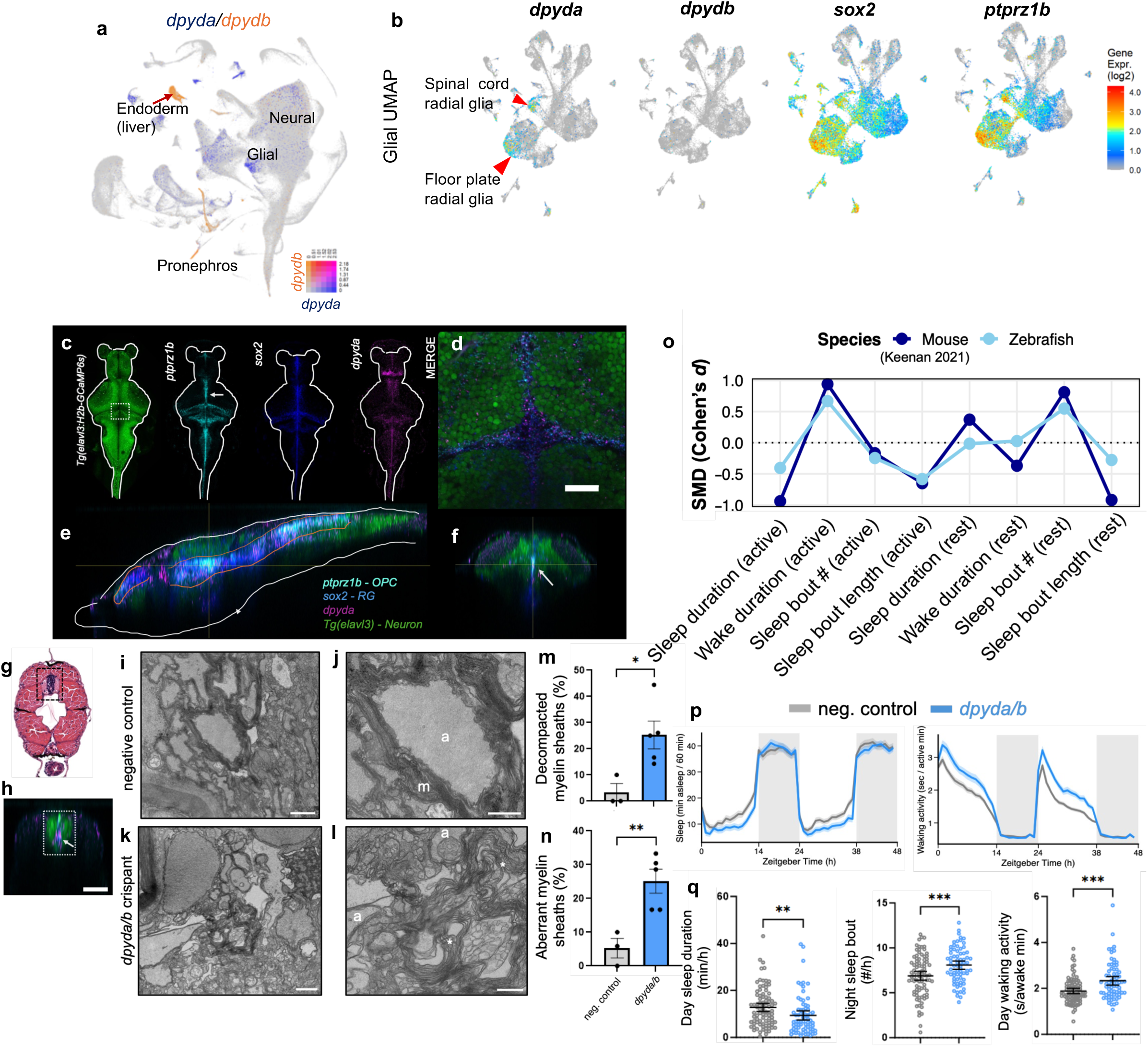
*dpyda/b* loss affects sleep and myelination in zebrafish similar to mouse. **a.** UMAP plots from Daniocell showing expression of both *dpyda* (blue) and *dpydb* (orange) across all cell types. Pink cells indicate expression of both. **b.** UMAP plots from Daniocell showing *dpydb, dpyda, sox2,* and *ptprz1b* expression in zebrafish glial cell types. **c.** Maximum intensity projections from confocal microscopy showing RNA localization by *in situ* HCR in whole-mount 6 dpf zebrafish for glial markers, *sox2* (radial glia-RG) *and ptprz1b* (oligodendrocyte precursor cells-OPC) along with *dpyda.* HCR was performed on *Tg*(*elavl3*:*H2B-GCaMP6s*) larvae labeling neuronal nuclei. **d.** Higher magnification view of ventricular domain indicated by the white box in c. scale bar is 25 µm. **e-f.** orthogonal projections of c showing colocalization along the ventricles (orange outline). White arrows in c and f indicate the z-slice in e. **g.** Transverse section of the zebrafish tail from bio-atlas.psu.edu showing approximate slice for TEM imaging. Spinal cord is outlined by the dashed box. **h.** Transverse projection of the approximate location displayed in g showing expression of the markers in c. White box matches black box in g. White Arrow indicates *dpyda* expression (magenta) in the ventral spinal cord. Scale bar = 50 µm. Representative TEM images taken from 8 dpf control (**i** and **j**) and *dpyda/b* crispants (**k** and **l**) showing myelinated axons of the ventral spinal cord (a = axon, m = myelin sheath). Examples of aberrant myelination marked with asterisks in l. **m.** Quantification (mean and SEM) of decompacted myelin sheaths shown as a percentage of total axons per fish. **n.** Quantification (mean and SEM) of aberrant myelin sheaths shown as a percentage of total axons per fish. n = 3 controls and 5 *dpyda/b* crispants. \**P*< 0.05, \*\**P*<0.01 represents unpaired student’s t-test. Scale bars in i and k = 1 µm and j and l = 500 nm. **o.** Standardized mean difference (SMD) calculated as Cohen’s *d* comparing zebrafish and mouse behaviors. Mouse data are from *Keenan et al. 2021*^8^. Active phase is relative to species as described in **Methods**. **p.** 48-hour trace showing mean and SEM of sleep duration and waking activity in *dpyda/b* crispants (n = 72) vs. negative control siblings (n = 88) pooled from two independent experiments. **q.** Quantification (mean and 95% CI) of selected measures in p. \*\**P*<0.01, \*\*\**P*<0.001 by Mann Whitney test for day sleep duration and day waking activity; unpaired student’s t-test for night sleep bout number.

Since radial glia are critical to the regenerative capacity in the zebrafish and serve as progenitors to oligodendrocytes^48,52^, we hypothesized that loss of the *DPYD* orthologs in zebrafish would impair myelination, as already observed in the mouse. To test this, we used CRISPR-Cas9 to mutate the conserved functional domains of both copies of *dpyd*, thereby generating loss-of-function mutations in both genes to prevent potential compensation^36,37,53,54^ (**Extended Data Fig. 3a-c**). CRISPR mutants are hereafter referred to as “*dpyda/b* crispants”. Similar to our studies in mice, we performed TEM on zebrafish axons to identify changes to myelin. Transverse sections of the distal spinal cord that contain prominent myelinated axons were collected from *dpyda/b* crispants and sibling controls injected with a non-targeting gRNA (**Fig. g-l**). Our *in situ* HCR data indicated *dpyda* is expressed in the spinal cord (**Fig. 2h**), and public single cell RNA-seq data show *dpyda* is expressed in radial glia and oligodendrocyte progenitors of the spinal cord (**Fig. 2b**). We noticed many decompacted myelin sheaths in *dpyda/b* crispants (**Extended Data Fig. 4**); thus, we assessed myelin compaction and appearance^55,56^ rather than g-ratio, which can be skewed due to myelin decompaction. While myelin was tightly packed in sibling controls with very few aberrant myelin membranes, *dpyda/b* crispants had significantly more membranes showing myelin decompaction (22% more decompacted sheaths, *P* = 0.03) (**Fig. 2m and Extended Data Fig. 4)** and aberrant myelin features (e.g. membrane vacuolization and irregular wrapping) (20% more aberrant sheaths, *P* = 0.009) (**Fig. 2n and Extended Data Fig. 4**). Together with our results from mice, we conclude that *DPYD* orthologs serve a conserved role in maintaining myelin across species.

Given one zebrafish ortholog to *DPYD* is expressed in glial progenitors and its loss affects myelination similar to mice, we tested whether *dpyda/b* crispants would produce a sleep phenotype consistent with that observed in *Dpyd* knockout mice. To directly compare the zebrafish phenotype to that we previously measured by electroencephalography (EEG) in *Dpyd* KO mice^8^, we compared effect sizes (Cohen’s *d*) for each equivalent measure. Across multiple sleep and wake measures, *dpyda/b* crispant zebrafish phenocopy *Dpyd* KO mice (**Fig. 2o**), even though in different light /dark phases, since zebrafish are nocturnal sleepers unlike mice. The significant phenotypes observed in mouse were characterized by reduced sleep during the active phase (lights off) and fragmented sleep during the inactive phase (lights on)^8^. This phenotype was also captured in our zebrafish *dpyda/b* crispants, where *dpyda/b* crispants slept an average of 47.1 minutes less during the active phase (lights on: *P* = 0.001) and had more fragmented sleep during the inactive phase (lights off: +11.9 sleep bouts/night, *P* = 0.0007) (**Fig. 2p-q and Extended Data Fig. 3**). *dpyda/b* crispants were hyperactive during their waking episodes (**Fig. 2p-q**) and had an intact arousal response to dark transitions (**Extended Data Fig. 3d-e**). These results were confirmed in two independent experiments (**Extended Data Fig. 3g**), showing high reproducibility of the crispant phenotype that recapitulates sleep/wake behaviors measured using EEG in mice.

Together, these results show *DPYD* orthologs in fish and mouse are conserved in expression and function similarly to regulate sleep.

### Manipulation of myelin affects sleep

We next asked whether direct manipulation of myelin could affect sleep and explain the phenotype observed following *DPYD* ortholog loss. Zebrafish hypomyelinating and hypermyelinating mutants have been well characterized; therefore, we used CRISPR-Cas9 to mutate two known myelin modifiers that have direct and opposing effects on myelin^32,57^(**Fig. 3a**). Guide RNAs were designed to disrupt critical functional domains predicted to result in protein-truncating, loss-of-function mutations^54^. First, we mutated the conserved transcription factor required for oligodendrocyte differentiation and myelination, *myrf*^32,58^ (**Fig. 3a and Extended Data Fig. 5**). Loss of *myrf* in zebrafish has been previously shown to result in fewer mature oligodendrocytes and thinner myelin^32^. *myrf* crispants slept on average 30.3 minutes less per light period than controls (*P* = 0.04), with no effect on daytime waking activity (*P* = 0.42) (**Fig. 3b-c**). Thus, *myrf* crispants produce a sleep loss phenotype similar to that of *dpyda/b* crispant zebrafish, indicating loss of oligodendrocyte myelination is sufficient to reduce sleep duration. Nighttime sleep was also reduced on average 56.6 minutes per dark period (*P* = 0.0015) with a concomitant increase in nighttime activity (77.5 seconds per dark period, *P* = 0.0019) following *myrf* loss (**Fig. 3b. and Extended Data Fig. 5**). The dark-transition arousal response was not impacted by *myrf* mutation (**Extended Data Fig. 5d-e**).

**Figure 3.**
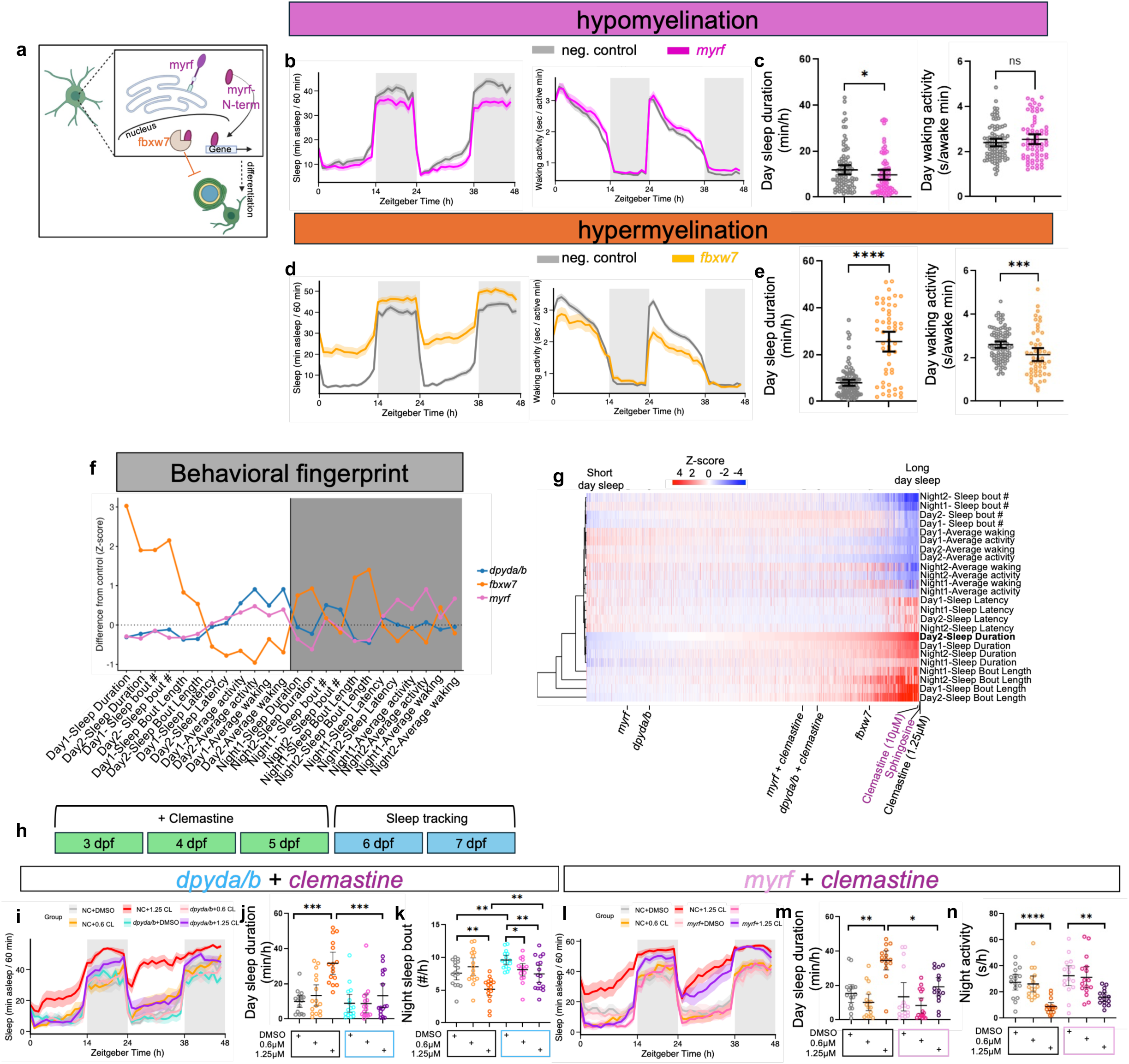
Genetic and pharmacological manipulation of myelin alters sleep. **a.** Schematic of *myrf*-*fbxw7* interaction adapted from *Nakayama et. al. 2018*^61^. **b.** 48-hour trace showing mean and SEM of sleep duration and waking activity in *myrf* crispants (n = 68) vs negative controls (n = 86) from two pooled experiments. **c.** Quantification of selected measures in b. **d**. 48-hour trace showing mean and SEM of sleep duration and waking activity in *fbxw7* crispants (n = 53) vs negative controls (n = 95) from two pooled experiments. **e.** Quantification of selected measures in d. **f.** Behavioral fingerprint plotting z-score difference from negative controls. Data are representative of two separate replicates for each experiment. Z-score is derived from the average z-score of the two replicates. **g.** Cluster plot of behavioral fingerprints for compounds from *Rihel et al. 2010*^63^ calculated as z-score deviation from controls. Data are sorted by Day 2 sleep duration. Compounds labeled in pink were tested by *Rihel et. al. 2010*^63^. Clemastine 1.25 µM was used in the current study. **h.** Schematic showing clemastine drug assay. **i.** 48-hour sleep trace showing mean and SEM for each group. CL = clemastine. Quantification of day sleep duration (**j**) and night sleep bout number (**k**) in negative controls (outlined by black box) and *dpyda/b* crispants (outlined by blue box) given 0.6 or 1.25 µM clemastine or DMSO. **l.** 48-hour sleep trace showing mean and SEM for each group. Quantification of day sleep duration (**m**) and night activity (**n**) in negative controls (outlined by black box) and *myrf* crispants (outlined by pink box) given 0.6 or 1.25 µM clemastine or DMSO. n = 15-16/ group in clemastine assays. ns = not significant \**P*<0.05, \*\**P*<0.01, \*\*\**P*<0.001, \*\*\*\**P*<0.0001 as determined by Mann-Whitney (waking activity and day sleep) or unpaired student’s t-test (nighttime sleep bout number) in c, and e, and Kruskal-Wallis followed by two-stage linear step-up procedure in **j**, **k**, **m**, and **n**. Quantification plots show mean and 95% CI. Measures were chosen based on those that differed significantly in sleep assays without drug administration.

To next test whether increasing myelination would have an opposite effect on sleep, we used CRISPR to disrupt *fbxw7* (**Extended Data Fig. 6**), which encodes a subunit of the E3 ubiquitin ligase that binds and degrades *myrf*^57,59–61^ (**Fig. 3a**). Previous work showed that loss of *Fbxw7* in mice and zebrafish results in an upregulation of *Myrf* that triggers accelerated oligodendrocyte differentiation and excess myelination^57^, which we hypothesized would have an opposite effect on sleep compared to *myrf* or *dpyda/b* loss. Indeed, *fbxw7* crispants showed an increase in daytime sleep duration (+246.2 minutes per 14-h light period, *P* < 0.0001) and reduced daytime waking activity (*P* = 0.0008). In a multivariable linear regression model controlling for daytime waking activity, the *fbxw7* genotype still maintained a robust effect on daytime hourly sleep duration (*ß* = 13.7 mins/hour, SE = 1.3, *P* < 0.001; overall model F_(2, 145)_ = 164.1, *R*^2^ = 0.69, *P* < 0.001), indicating reduced waking activity does not fully explain the longer sleep duration. Additionally, the dark-transition response was preserved demonstrating intact arousal systems in *fbxw7* crispants (**Fig. 3d-e and Extended Data Fig. 6**). This demonstrates *fbxw7* loss, a model of hypermyelination, increases sleep duration.

We generated a “behavioral fingerprint” for each mutant by calculating the z-score deviation for different aspects of the phenotype relative to negative control-injected siblings^62,63^ to compare the sleep and activity profiles between hypermyelinating and hypomyelinating mutants. Almost every behavioral measure showed an opposite effect between *myrf* and *fbxw7* crispants (**Fig. 3f**), confirming the known regulatory interaction between these two genes^57,60,61^. Moreover, *dpyda/b* crispants largely phenocopied the *myrf* crispants across behavioral measures, although each had distinct effects on day and night measures (**Fig. 3f**). Our results support the hypothesis that *dpyd* influences sleep through its effect on myelin.

To further rule out the possibility that disruption of peripheral pyrimidine catabolism to *β*-alanine contributes to the sleep loss in *dpyda/b* crispants, we generated zebrafish with LOF mutations in the genes encoding the two other pyrimidine catabolic enzymes *dpys* and *upb1*, which are primarily expressed in the liver (**Extended Data Fig. 7a**). Mutation of *dpys* did not affect sleep (**Extended data Fig. 7b-d**), and *upb1* crispants displayed an opposite phenotype to *dpyda/b* crispants with reduced activity along with increased day sleep (**Extended Data Fig. 7e-g**). *upb1* crispants were less responsive to light touch during early development and after accounting for daytime waking activity, *upb1* crispants showed no independent change in daytime hourly sleep compared to controls (*ß* = 1.4 mins/hour, SE = 1.4, *P* = 0.31), demonstrating that the observed sleep variations in *upb1* crispants are secondary to alterations in gross motor activity. Collectively, our data do not support a role for peripheral pyrimidine catabolic end products as the source of the sleep phenotype following *dpyda/b* loss.

We next tested whether pharmacological manipulation of myelin could alter sleep. To identify plausible candidate compounds that affect sleep and myelin, we aligned our behavioral data with a previously published large-scale small molecule screen^62,63^ that assessed multi-day effects on larval zebrafish sleep and activity, creating behavioral fingerprints (similar to Figure 3f) for each compound. Aligning mutant sleep-wake behaviors with these behavioral fingerprints has previously been successful in predicting pharmacological agents that can rescue or synergize with mutant phenotypes^37,64^. Data were sorted by daytime sleep duration to identify compounds with opposing sleep patterns to hypomyelinating *dpyda/b* and *myrf* crispants, which clustered with compounds that reduced sleep (**Fig. 3g**). Sphingosine and clemastine, which promote myelination^65–69^, were two of the strongest daytime sleep-promoting compounds (**Fig. 3g**), providing independent support that enhancing myelination is sleep promoting. Clemastine is an FDA-approved compound that has been shown to promote myelination across species by increasing oligodendrocyte precursor proliferation and differentiation^65–72^. Given this, we tested whether clemastine could restore sleep in our hypomyelinating mutants.

We first tested multiple doses of clemastine alongside the potent sleep-promoting compound, melatonin as a positive control. Drugs were applied via bath application to wild-type zebrafish to determine a dose of clemastine that would elicit a sleep-inducing effect similar to that observed previously^63^ while minimizing toxicity, as indicated by abnormal growth or death. Six doses of clemastine (0.625, 1.25, 2.5, 5, 10 and 20 µM) were tested with limited toxicity at 0.625 µM and 1.25 µM doses applied from days 3-5 post fertilization (**Fig. 3h and Extended Data Fig. 8**). Doses as low as 500 nM have been shown to induce oligodendrocyte proliferation and myelination in zebrafish larvae^69,71^. Drug application was stopped prior to sleep analysis on days 6 and 7 post fertilization, though residual drug remained in the wells. Clemastine doses above 2.5 µM were either toxic or completely abolished activity (**Extended Data Fig. 8**). A potential caveat to using clemastine is that at certain doses it is a histamine H1 receptor antagonist, which would be expected to induce sleep^73,74^. Zebrafish exhibit a histamine-dependent arousal response to the light-to-dark transition^73^. Thus, we looked for a dose that induces sleep but preserves this dark-transition response (**Extended Data Fig. 8**), indicating histamine antagonism is not the primary driver of the behavioral quiescence. All doses 2.5 µM and above abolished the dark-transition response; however, 1.25 µM clemastine increased daytime sleep in wild-type larvae from 6-7 dpf with little impact on nighttime sleep and no effect on the dark-transition response (**Extended Data Fig. 8d-g**). Both lowest doses of clemastine (0.625 and 1.25 µM) preserved the histamine H1 receptor-dependent dark-transition hyperlocomotion response (**Extended Data Fig. 8f-g**), suggesting the sleep-inducing effect of low dose clemastine is not due to H1 antagonism.

Based on these dose-response analyses in wild-type fish, we selected the two lowest doses to apply to *dpyda/b* and *myrf* crispants and found that 1.25 µM restored sleep and activity parameters in both hypomyelinating mutants to baseline levels found in controls given DMSO. However, hypomyelinating mutants given clemastine showed an attenuated response compared to controls given clemastine (**Fig. 3i-n and Extended Data Fig. 9**), indicating *dpyd* and *myrf* are important for the full sleep-promoting effect of clemastine, likely because clemastine induces transcription of genes required for oligodendrocyte myelination such as *Myrf*^72^. Comparison of the behavioral fingerprints following drug administration demonstrated *dpyda/b* and *myrf* crispants given clemastine normalized to a mid-range sleep duration (**Fig. 3g**). Importantly, *dpyda/b* and *myrf* crispants showed no difference in the histamine-dependent dark-transition response compared to their respective controls (**Extended Data Fig. 9e and j**), suggesting that differences in histaminergic responsiveness do not explain the *dpyd/myrf*-dependent sleep phenotype. Additionally, *dpyda/b* crispants responded to 1 µM melatonin to the same degree as controls (**Extended Data Fig. 10**), indicating *dpyda/b* crispants are capable of sleep induction via other mechanisms.

Collectively, our data indicate hypomyelination and hypermyelination produce opposing sleep phenotypes, and the promyelinating compound clemastine can ameliorate sleep deficits in hypomyelinating mutants, including *dpyda/b* crispants.

### *Drosophila su(r)* alters sleep and glial lipids

Thus far, our experiments support the hypothesis that DPYD affects sleep by altering oligodendrocyte myelination, raising the question of how myelin could directly influence sleep. Myelin could act through its established insulating role in enhancing axonal conduction velocity and/or through its capacity to provide metabolic substrate to wake-active neurons as a lipid reservoir^75–77^. To distinguish between these possible mechanisms, we turned to *Drosophila*, which lack wrapping myelin in the CNS entirely but possess a conserved ortholog to *DPYD*, *su(r)*. Reanalysis of published single-cell RNA-sequencing data from the Fly Cell Atlas^78^ indicate that *su(r)* is expressed predominantly in glia, particularly ensheathing glia and peri/subperineurial glia (**Fig. 4a**). *Drosophila* glia do not myelinate axons in the CNS, but they enwrap neuronal cell bodies and provide metabolic support through lipid storage organelles called lipid droplets^79,80^, which have been proposed as an evolutionary precursor to myelin^76,81^. Mobilization of glial lipid stores from lipid droplets has been shown to be sleep-promoting in *Drosophila*^82^; therefore, altering this lipid storage is expected to affect sleep. Thus, studies in *Drosophila* allow us to ask whether a DPYD-dependent, lipid-storage-based mechanism could support sleep in the absence of wrapping myelin. If DPYD’s effect on sleep depends specifically on the insulating properties of wrapping myelin, loss of the invertebrate ortholog should not affect sleep; if DPYD instead acts through a glial lipid-storage function to support brain metabolism, we would expect a sleep phenotype in flies despite the absence of myelin.

**Figure 4.**
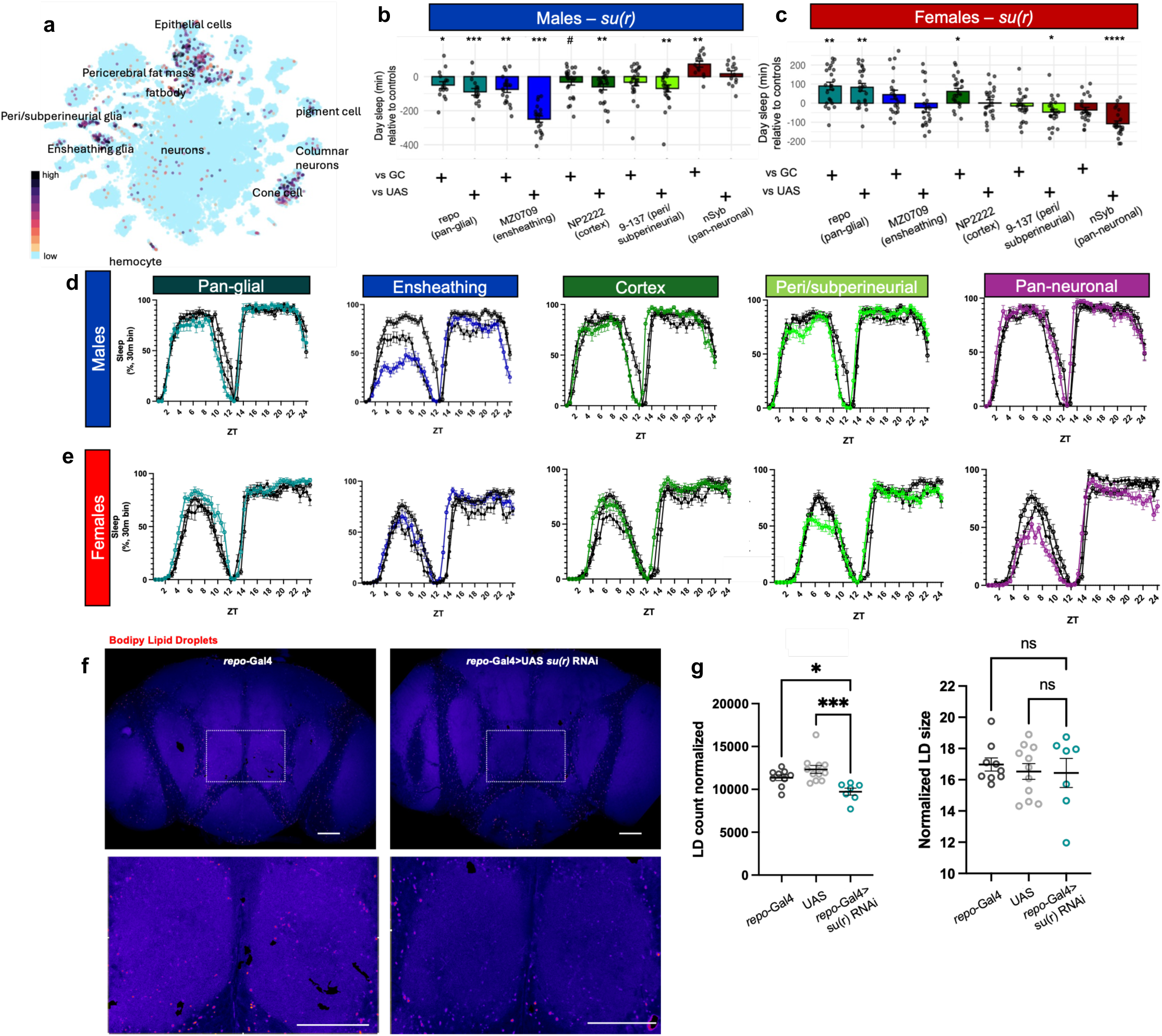
*Drosophila* ortholog to *DPYD*, *su(r)*, is expressed in glia and influences sleep and lipid droplets. **a.** UMAP plot showing *su(r)* expression in different cell types from *Drosophila* head. Data are from flycellatlas.org. Day sleep duration plotted as minutes lost or gained relative to the GAL4-only control (GC) or UAS-only control for male flies (**b**) and female flies (**c**). Points represent individual flies. Sleep traces plotted as percent time asleep per 30-minute bin across 24 hours for males (**d**) and females (**e**). Data are plotted as mean and SEM for GAL4>UAS *su(r)* RNAi (colored line) and GAL4, or UAS-only controls (black and gray lines). **f.** Representative images of fly brains stained using BODIPY^493/503^ to label lipid droplets (LD). Scale bar = 50 µM. Bottom image is inset of dashed box in top. **g.** Quantification (mean and SEM) of LD count and size. ^#^*P*<0.1, \**P*<0.05, \*\**P*<0.01, \*\*\**P*<0.001, \*\*\*\**P*<0.0001 determined by one-way ANOVA with Holm-Sidak’s multiple comparisons test. Sample n as follows for male: *repo*>su(r) RNAi = 16, *repo* GAL4/+ = 21; MZ0709>su(r) RNAi = 21, MZ0709 GAL4/+ = 20; NP2222>su(r) RNAi = 24, NP2222 GAL4/+ = 24; 9-137>su(r) RNAi = 23, 9-137 GAL4/+ = 24; *nSyb*>su(r) RNAi = 14, *nSyb* GAL4/+ = 26; +/UAS su(r) = 24 (all groups). n for females: *repo*>su(r) RNAi = 24, *repo* GAL4/+ = 20; MZ0709>su(r) RNAi = 21, MZ0709 GAL4/+ = 19; NP2222>su(r) RNAi = 24, NP2222 GAL4/+ = 25; 9-137>su(r) RNAi = 24, 9-137 GAL4/+ = 24; *nSyb*>su(r) RNAi = 22, *nSyb* GAL4/+ = 24; +/UAS su(r) = 24 (all groups).

We first performed pan-glial knockdown of *su(r)* using *repo*-GAL4 in male flies, which selectively reduced sleep during the day (-48.4 minutes vs. *repo*-GAL4/+, *P* = 0.03; -89.5 minutes +/UAS *su(r)* RNAi, *P* = 0.0002) (**Fig. 4b**) driven by a reduction in sleep bout length and increased activity (**Extended Data Fig. 11**). Effect size comparison across mouse, zebrafish, and *Drosophila* show this effect on sleep and wake during the animal-specific active phase is common across all species (**Extended Data Fig. 11a**). Knockdown using drivers targeting specific glial subtypes—cortex glia (NP2222), peri/subperineurial glia (9-137), and ensheathing glia (MZ0709)—showed that *su(r)* regulates sleep primarily via ensheathing glia (**Fig. 4b, d**), consistent with its high expression in this cell type. Neuronal knockdown using *nSyb*-GAL4 did not significantly affect sleep (**Fig 4b, d, and Extended Data Fig. 11**). These results demonstrate that *su(r)* loss is sufficient to reduce sleep in the absence of myelin, indicating that DPYD can act through a glia-intrinsic mechanism that does not require myelin wrapping.

We did not observe sex differences in mice lacking *Dpyd*^8^, and larval zebrafish are not sexually differentiated^83^. However, *Drosophila* have previously been shown to have significant sex differences in daytime sleep duration^84^, and *su(r)* is on the X chromosome. Therefore, we next asked whether this glial mechanism shows sex-specific regulation in *Drosophila*. Cell-specific knockdown in female flies revealed effects different to males: pan-glial (*repo*) knockdown in females significantly *increased* daytime sleep duration (+89.7 minutes vs. *repo*-GAL4/+, *P* = 0.008; +83.3 minutes vs. +/UAS *su(r)* RNAi, *P* = 0.008), (**Fig. 4c, e, and Extended Data Fig. 11**). Females were generally more resilient to *su(r)* loss than males, in that knockdown in all other glial cell types and neurons did not significantly impact daytime sleep compared to both parental controls (**Fig. 4c, e, and Extended Data Fig. 11**), which may represent compensation^85^. Across both sexes, significant effects were primarily restricted to day sleep, consistent with the previously reported peak expression of this gene in *Drosophila* heads^86^.

To test whether this glial mechanism reflects a lipid-storage function, we asked whether *su(r)* loss affects glial lipid droplet formation. *DPYD* has been shown to promote lipid droplet formation in human hepatocytes^16^; thus, a similar function in glia may exist. We performed pan-glial knockdown of *su(r)* using *repo*-GAL4 to prevent compensation from unique glial subtypes and found this significantly reduced the number of lipid droplets in the *Drosophila* brain (*P* = 0.02 vs. *repo*-GAL4/+, *P* = 0.0006 vs. +/UAS *su(r)* RNAi) (**Fig. 4f-g**). Together, these results indicate that *su(r)* supports both sleep and glial lipid droplet formation, consistent with a model in which this *DPYD* ortholog alters sleep through a glial lipid-storage mechanism that may serve a similar metabolic role to vertebrate myelin.

### Genome wide association studies (GWAS) of sleep implicate *DPYD* in oligodendrocytes

Our cross-species analysis indicates orthologs to *DPYD* exert their effects on sleep through glia, specifically of the oligodendrocyte lineage. To determine whether this implied mechanism is supported by the genetic association for sleep duration reported at the human *DPYD* locus, we intersected the GWAS data for sleep duration with public epigenomic data collected in oligodendrocytes^87^. Epigenetic signatures are cell-type specific and reveal regulatory elements required for gene localization and cell-specific function.

We obtained the GWAS sentinel variants for accelerometer-derived sleep duration^2^ and correlated proxy variants for these sleep association signals using LDproxy^88^ (**Fig. 5a**) to identify potential causal variants at the *DPYD-MIR137* GWAS locus with regulatory potential. Variant rs2660304 (*r*^2^ with sentinel variant rs2660302 = 0.69) was selected as a candidate causal proxy as this variant had the highest FORGEdb^89^ score, indicating significant regulatory potential. Importantly, eQTL data from blood^90^ show rs2660304 (G), which is associated with shorter sleep duration^2^, is also associated with reduced expression of *DPYD* (z-score = -10.1, *P* = 7.64 × 10^-24^); thus, our LOF models are relevant to this human GWAS variant. rs2660304 falls in a region marked by accessible chromatin (ATAC-seq peak) that shows high H3K27ac and H3K4me3 in oligodendrocytes indicating an active promoter^91,92^ (**Fig. 5b-c**). This variant resides in the *MIR137* host gene, but chromatin interaction data from oligodendrocytes^87^ show rs2660304 forms a physical chromatin contact with the accessible promoter of *DPYD* and a distal intronic enhancer, similarly marked by accessible chromatin in oligodendrocytes (**Fig. 5b-c**). Notably, this interaction is absent in neurons and microglia^87^ (**Extended Data Fig. 12**), suggesting the variant may regulate the promoter of *DPYD* specifically in oligodendrocytes. The region harboring rs2660304 shows high conservation despite residing within a non-protein-coding region; therefore, we leveraged public cis-regulatory atlases to examine the analogous region across mouse^93^ (**Fig. 5d**), zebrafish^94^ (**Fig. 5e**), and *Drosophila*^95^ (**Fig. 5f**). This regulatory chromatin signature is preserved across all species. The promoter of *DPYD* orthologs is marked by accessible chromatin (ATAC-seq peaks) in mouse OPCs, zebrafish radial glia, and *Drosophila* ensheathing glia (**Fig. 5c-e**), indicating this gene is accessible to transcription in the glial cell types in which they are most highly expressed in each species. *MIR137* and *DPYD* are syntenically conserved in vertebrates, and the same chromatin architecture is observed between *MIR137* and *DPYD* in both mouse and zebrafish, suggesting that the sleep duration GWAS variant rs2660304 resides in a regulatory element that is likely conserved.

**Figure 5.**
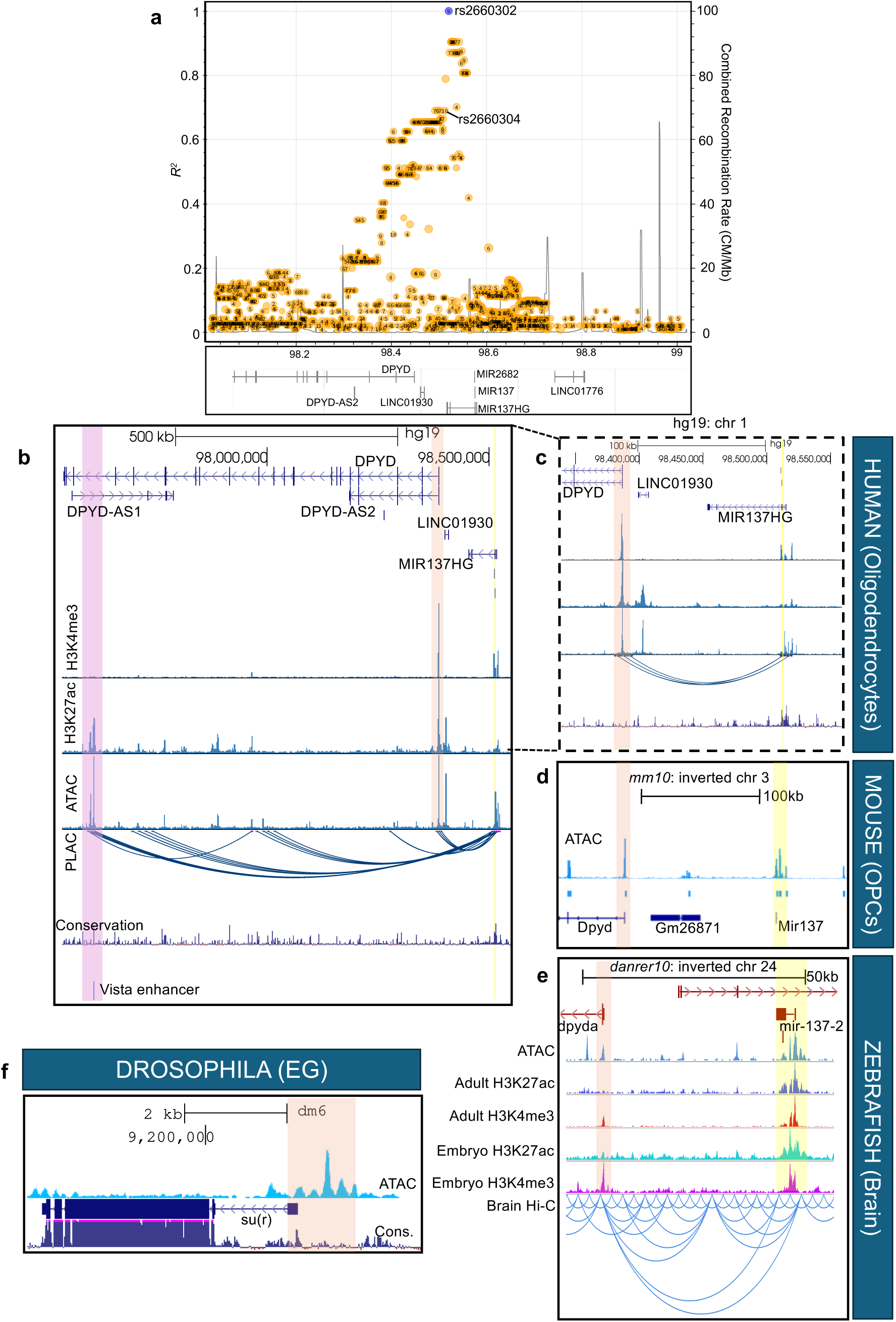
Human GWAS variants for sleep duration implicate a role for *DPYD* in oligodendrocytes. **a.** Locus zoom plot from LDLink showing *r*^2^ correlation for index variant rs2660302 for accelerometer-derived sleep duration reported by *Jones et. al., 2019*^2^. **b-c.** UCSC Genome Browser plot showing the human (hg19) *DPYD-MIR137* locus. Correlated proxy variant (*r*^2^ = 0.69), rs2660304 (yellow line), falls within an internal promoter of the *MIR137* host gene (*MIR137HG*). This variant resides in a region showing active histone marks in oligodendrocytes and forms physical chromatin contacts with the promoter of *DPYD* (orange highlight) and a distal intronic vista-confirmed enhancer (purple highlight) in oligodendrocytes. The *DPYD* promoter and distal intronic enhancer show histone marks indicating an accessible active promoter (H3K4me3 + H3K27ac + ATAC) and enhancer (H3K27ac + ATAC), respectively. Chromatin H3K27ac, H3K4me3, ATAC-seq and PLAC-seq data are from *Nott et al., 2019*^87^. Conserv. = vertebrate conservation. **d.** The analogous region (inverted for comparison to human) for *Dpyd* and *Mir137* against the mouse genome (mm10), showing ATAC-seq data from OPCs^93^. **e.** The analogous region (inverted) for *dpyda* and *dre-mir137-2* against the zebrafish genome (danrer10). Chromatin data are from *Yang et. al. 2020*^94^ showing marks of active promoters (H3K27ac + H3K4me3 + ATAC) and enhancers (H3K27ac + ATAC). Hi-C chromatin interaction data from zebrafish brain show interaction between *dre-mir137-2* and the *dpyda* promoter as in human. Single cell ATAC-seq data in zebrafish are from the sox9 cluster representing radial glia**. f.** The analogous region against the *Drosophila* genome (dm6) showing *su(r)* along with ATAC-seq data from ensheathing glia^95^. The insect conservation (cons.) track is shown below. In b-e, the orange highlight marks the *DPYD* promoter region across species while the yellow highlight marks the *MIR137* region that harbors the GWAS association in humans. *MIR137* is conserved in *Drosophila* but is on a different chromosome.

These findings importantly bridge the human GWAS variant for sleep duration to active regulation of *DPYD* in oligodendrocytes, implicating a cellular mechanism underlying this genetic association to sleep duration in humans and across species.

## Discussion

Our cross-species analysis identified DPYD as a conserved regulator of sleep that acts within specialized glial populations—ensheathing glia in flies, radial glia in fish, oligodendrocyte progenitors in mouse, and mature oligodendrocytes in human. This evolutionary pattern of expression is consistent with the finding that metabolite conversion enzymes, like DPYD, evolve with cell specialization through cell type subfunctionalization^96^. This conservation, spanning myelinating and non-myelinating nervous systems, points to an ancestral glial function for DPYD that has been retained even as these cell types diversified and is supported by our finding that DPYD orthologs affect glial lipids across species, though, through different properties. In each model system we studied, loss of the *DPYD* ortholog showed a significant effect on sleep duration during the species’ active phase, consistent with the rhythmic expression of this enzyme reported previously^86,97,98^ and with sleep phenotypes observed following loss of other glial regulators^99,100^.

A central question raised by our results is: what feature of glial biology links *DPYD* to sleep across such divergent nervous systems? We show that *DPYD* loss reduces brain lipids stored as myelin in vertebrates and as lipid droplets in *Drosophila*, and that the *Drosophila* ortholog *su(r)* regulates sleep even in the complete absence of myelin. This indicates that *DPYD* can act through a glial lipid-storage mechanism that may pre-date wrapping myelin’s functions, which is consistent with the hypothesis that glial lipid droplets represent a functional evolutionary precursor to myelin^81,101,102^, though direct evidence for this relationship remains to be established. Mechanistically, DPYD inhibition in hepatocytes downregulates cholesterol biosynthesis genes including *HMGCS1*^16^, and data from the International Mouse Phenotyping Consortium (https://www.mousephenotype.org/) show *Dpyd* knockout mice have reduced cholesterol and triglycerides, suggesting DPYD loss may impair myelin biogenesis by limiting substrate availability for lipid biosynthesis or membrane stability^103^. Existing literature also implicates lipids directly as sleep-promoting signaling molecules^104–107^, raising the possibility that lipid availability itself, independent of storage structure, is permissive to sleep.

Beyond the mechanism related to *DPYD*, our zebrafish experiments address an open question in the field: whether the relationship between sleep and myelination is bidirectional. Sleep is known to promote myelin gene expression, and sleep restriction impairs myelination^24,27,30,108^, but whether myelin content reciprocally influences sleep had not been directly tested. We show that reducing myelin decreases sleep while enhancing myelin is permissive to sleep, indicating oligodendrocytes are not merely responders to sleep loss but active regulators of sleep. A recent study knocking out Bmal1 in mouse oligodendrocyte precursors led to myelin loss and fragmented sleep, supporting a direct role for these cells^100^. However, Bmal1 loss may influence broader transcriptional coordination that contributes to sleep. Collectively, our data support a bidirectional relationship between sleep and myelination and suggest that an optimal level of myelin is important for the maintenance of sleep duration. These findings have direct relevance to demyelinating and neurodegenerative diseases. For example, sleep is disrupted in multiple sclerosis^109^ and Alzheimer’s disease^110^, and degrades with age alongside myelin loss^111,112^. Our findings raise the possibility that sleep disruption is an early signal of myelin pathology, which could provide an avenue for early intervention in these disorders.

Our results highlight the therapeutic potential: clemastine, an FDA-approved remyelinating compound already in clinical trials for demyelinating disease^65,66^, rescued aberrant sleep and activity in our hypomyelinating zebrafish mutants. The behavioral fingerprinting approach, aligning mutant phenotypes against a previously published compound screen^62,63^, offers a scalable strategy for identifying additional compounds that jointly target sleep and myelin deficits. Pharmacological and genetic manipulation of myelin-related genes may impact additional targets, and we cannot rule out all impacts on other neuromodulatory systems; however, our combinatorial approach supports the concept that increasing myelin increases sleep and reducing myelin shortens sleep.

DPYD’s role in lipid metabolism may help explain its pleiotropic genetic—GWAS— associations with metabolic^113,114^, cognitive^115^, and neuropsychiatric traits^116^, and argues for considering CNS effects when using DPYD inhibitors such as those prescribed alongside 5-fluorouracil chemotherapy. Evidence of CNS involvement is indicated by rare case reports of delayed myelination and diverse neurological phenotypes in patients with DPYD loss-of-function variants^117,118^ and by reports of DPYD upregulation in glia during neurodegeneration^119^, ageing^120^, and MS^121^. The sleep-duration GWAS variant at this locus is also in linkage disequilibrium with a major schizophrenia risk association^116^, a disorder independently marked by myelin abnormalities^122^, suggesting this regulatory element may warrant deeper investigation across a broader set of brain-related traits.

Several questions remain including how DPYD specifically affects lipid synthesis and/or storage in the brain and whether its effects extend beyond myelin. It is curious that DPYD is highly expressed in cell types, including glia, that do not express the other enzymes in the pyrimidine catabolism pathway. This suggests the immediate metabolites of DPYD, dihydrouracil and dihydrothymine, may serve a unique role in these cells, for example as signaling molecules^123^. How myelin and brain lipids in general contribute to sleep is also an open question. *Drosophila* studies indicate beta oxidation of lipid stores is key to sleep regulation, but whether this is due to direct lipid signaling or restoration of energetic demands is still under investigation^77,82,124–127^. Whether myelin’s effect on sleep in vertebrates operates through this same lipid-storage function, through conduction-related properties, or both, was not directly tested here and remains an important question for future work. Additionally, further experiments are necessary to determine whether oligodendrocytes or their precursors serve independent functions in sleep regulation.

Using this integrated cross-species approach, we were well-placed to test distinct hypotheses. Each species offers a unique benefit: mouse allows for translational sleep and myelin characterization in a mammalian system, zebrafish facilitate genetic and pharmacological manipulation of myelination without severe developmental abnormalities, and the *Drosophila* GAL4-UAS system offers cell-specific interrogation. Altogether, our work identifies *DPYD* as a conserved regulator of glial lipid metabolism and sleep and provides direct evidence that myelin modification causally influences sleep.

## Methods

### Animal use

Mouse experiments were conducted in accordance with the Institutional Animal Care and Use Committee guidelines under the appropriate institutional approved protocol (805817) at the University of Pennsylvania. All experiments with zebrafish were conducted in accordance with the Institutional Animal Care and Use Committee guidelines under the approved protocols (806646 and IAC 21-001154) at the University of Pennsylvania and the Children’s Hospital of Philadelphia. Breeding pairs consisted of wild-type AB and TL (Tupfel Long-fin) strains for all sleep and genetic manipulation experiments. Fish for HCR imaging experiments were housed and bred in the UCL Zebrafish Facility under Project License PP6325955 awarded to Jason Rihel under the UK Animals (Scientific Procedures) Act 1986. Experimental procedures were in accordance with the UK Home Office Animals (Scientific Procedures) Act 1986. *Tg(elavl3:H2B-GCaMP6s)jf5Tg* homozygous for the *mitfa^w^*^2^ allele were used for HCR experiments. Fish were housed in standard conditions with 14-hour:10-hour light:dark cycle at 28.5°C, with lights on at 9am (ZT0) and lights off at 11pm (ZT14).

### Mouse sleep tracking

Mice used in this study have been previously characterized for sleep^8^ and are available through the Toronto Center for Phenogenomics International Mouse Phenotyping Consortium (C57BL/6N-Dpyd^tm1b(KOMP)Wtsi>^/Tcp; http://www.mousephenotype.org). Heterozygous breeding pairs were used to generate homozygous knockout mice and wild-type littermate controls. Mice were approximately 3 months of age at the time of study. Five males and one female were used given no sex differences were observed in our prior study^8^. Mice were individually housed in a sound-attenuated facility under a 12-hour:12-hour light:dark schedule. Temperature was regulated between 22-25°C. Humidity was 40-60%. Food and water (chlorinated tap water) were provided *ad libitum*. Mice were acclimated for at least one day prior to recording. We recorded activity patterns for 4 days by recording in 10-second epochs the number of times a mouse crossed electronic beams that intersected the mouse cage (Micromax, Accuscan Instruments, Inc.). Sleep was defined as 40 seconds or more of continuous inactivity. This definition has been validated against electroencephalography (EEG)^7^. In our current studies in mice, sleep was not a primary outcome as this has been previously assessed by EEG/ EMG recording in a larger cohort^8^.

### Liquid chromatography mass spectrometry

Adult male mice (Homozygous mutants [KO], heterozygous [HET], and wild-type [WT] controls) aged 3-8 months were sacrificed between ZT6-8, and brains were frozen on dry ice and stored at -80°C until dissection. Piriform cortices (3mm) were dissected and processed by the University of Pennsylvania Metabolomics Core. Spatial transcriptomic data from mouse brain (https://xenium.10xgenomics.com/) show *Dpyd* is highly expressed in the piriform cortex.

Polar metabolites were extracted from weighed mouse brain tissue using a modified Bligh–Dyer extraction. Briefly, tissue was homogenized in ice-cold 2:1 methanol:chloroform, followed by addition of water and chloroform to induce phase separation. Samples were centrifuged at 18,787 × g for 7 min at 4°C, and the upper aqueous phase containing polar metabolites was collected and dried under vacuum. Dried extracts were reconstituted in 1:1 acetonitrile:water and analyzed by hydrophilic-interaction ultra-performance liquid chromatography–tandem mass spectrometry (UPLC–MS/MS) using an XBridge BEH Amide column on a Waters ACQUITY H-Class UPLC coupled to a Waters Xevo triple-quadrupole mass spectrometer. Targeted metabolites were measured using multiple-reaction monitoring (MRM), with samples analyzed in randomized order and pooled quality-control samples included throughout the analytical run. Chromatographic peaks were identified and integrated using TargetLynx/MassLynx, and integrated ion counts were exported for subsequent processing. For this dataset, ion counts were normalized to the initial tissue weight and corrected for analytical drift across the run. Samples acquired after loss of instrument signal were excluded, and metabolites with a quality-control coefficient of variation >30% were removed; 14 of the 17 targeted metabolites passed this criterion and were retained for downstream analysis. The general extraction and LC–MS/MS workflow has been described previously^128,129^.

Prior to statistical testing, technical replicates for each animal were averaged to yield a single normalized intensity value. Differences in normalized metabolite intensities across the three genotype groups (WT, HET, and KO) were evaluated using the non-parametric Kruskal-Wallis test. For post-hoc comparisons between specific groups (WT vs. HET and WT vs. KO), pairwise Wilcoxon rank-sum tests (Mann-Whitney U tests) were conducted.

### Transmission electron microscopy

#### Mouse

Mice were sacrificed 6 hours into the light cycle (ZT 6) by cardiac perfusion using 10 mL of 1x PBS followed by 60 mL of 4% PFA. Brains were dissected and a 2 mm slice was taken from the medial corpus callosum isolated using Bregma coordinates, then placed into EM buffer of 2.5% glutaraldehyde, 2.0% paraformaldehyde in 0.1M sodium cacodylate buffer, pH7.4 and stored at 4℃ until processing. After subsequent buffer washes, the samples were post-fixed in 2.0% osmium tetroxide with 1.5% K_3_Fe(CN)_6_ for 1 hour at room temperature and rinsed in dH_2_O prior to *en bloc* staining with 2% uranyl acetate. After dehydration through a graded ethanol series, the tissue was infiltrated and embedded in EMbed-812 (Electron Microscopy Sciences, Fort Washington, PA). Thin sections were stained with uranyl acetate and SATO lead and examined with a Talos L120C electron microscope fitted with a Ceta 16M digital camera. Images were collected to capture axonal cross sections. Image acquisition was collected blinded to genotype. Images were analyzed using semi-automated quantification in FIJI with MyelTracer^130^ by two independent scorers blinded to genotype.

#### Zebrafish

Larval zebrafish were collected at 8 dpf for electron microscopic examination. Larvae were euthanized by rapid cooling on ice slurry, and eyes were removed for genotyping while bodies were placed into electron microscopy fixative solution of 2.5% glutaraldehyde, 2.0% paraformaldehyde in 0.1M sodium cacodylate buffer, pH7.4, overnight at 4℃. After subsequent buffer washes, the samples were post-fixed in 2.0% osmium tetroxide with 1.5% K_3_Fe(CN)_6_ for 1 hour at room temperature and rinsed in dH_2_O prior to *en bloc* staining with 2% uranyl acetate. After dehydration through a graded ethanol series, the tissue was infiltrated and embedded in EMbed-812 (Electron Microscopy Sciences, Fort Washington, PA). Thin sections were stained with uranyl acetate and SATO lead and examined with a Talos L120C electron microscope fitted with a Ceta 16M digital camera. Images were collected from the ventral spinal cord. Image acquisition was collected and analyzed blinded to genotype.

### Statistical Analysis of TEM images

#### Mouse

To evaluate the effect of genotype on g-ratio, we used linear mixed-effects models^30^ implemented in R with the *lme4* package, including a fixed effect for genotype and a random intercept for each animal. Significance of the genotype effect was determined by comparing the model with genotype to a null model without genotype using a likelihood ratio test. To determine whether genotype effects were modified by axon diameter, we performed an interaction test by including fixed effects of genotype, axon diameter, and their interaction in the mixed effects model. Likelihood ratio tests were used to compared nested models with and without the interaction terms to assess evidence for effect modification, using *lmerTest* in R.

#### Zebrafish

Aberrant myelin (multiple outfolds, separation from axon) and decompacted myelin (clear space between wraps) was defined visually in FIJI. Numbers of aberrantly myelinated axons or axons with decompacted myelin were totaled across images collected from each individual fish then divided by the total number of myelinated axons in the field of view for that fish. Unpaired student’s t-test was used to test group differences.

### CRISPR-Cas9 mutagenesis in zebrafish

CRISPR RNAs (crRNAs) were designed using the online tool Crispor (http://crispor.tefor.net/) as described previously^131^. crRNAs were designed to target all transcript isoforms annotated by Ensembl and a conserved protein domain critical to function (identified by DIOPT) to increase the likelihood of generating a non-functional protein^54^. Alt-R^TM^ crRNAs were ordered from IDT (https://www.idtdna.com/). AB/TL breeding pairs were set up overnight and embryos collected in embryonic growth media (E3 medium; 5mM NaCl, 0.17 mM KCl, 0.33 mM CaCl2, 0.33 mM MgSO4) the following morning shortly after lights-on. Pre-formed ribonucleoprotein (RNP) complexes containing the crRNA:tracrRNA (guideRNA, gRNA) and SpCas9 HiFi enzyme (IDT) were injected at the single-cell stage as previously described^36,37^. Injections of scrambled (non-targeting) gRNA and gene-specific gRNA were performed in embryos from the same breeding event (clutch). Embryos were left unperturbed for one day before being transported to fresh E3 media in Petri dishes (approximately 50 per dish).

To generate *dpyda/b* crispants, the conserved catalytic domain encoded by exon 4 was targeted in both *dpyda* and *dpydb* as this is the most upstream functionally conserved domain required for protein function (**Extended Data Fig. 3**). To generate *myrf* crispants, gRNAs were designed to target an upstream exon, exon 3, and the conserved NDT80 DNA binding domain encoded by exon 8 (**Extended Data Fig. 5**). To generate *fbxw7* crispants, a gRNA was designed to target exon 7, which encodes the conserved F-box domain required for interaction with the E3 ubiquitin ligase complex^57^ (**Extended Data Fig. 6**). To generate *upb1* crispants, a gRNA was designed to target exon 5, encoding the carbon-nitrogen hydrolase, and *dpys* crispants were generated with a gRNA targeting exon 5, which encodes the catalytic domain. crRNA sequences and PCR genotyping information is provided in **Supplementary Table 1**.

### DNA extraction and PCR for genotyping zebrafish

DNA extraction was performed per the manufacturer’s protocol (Quanta bio, Beverly, MA) immediately following completion of the sleep assay, as described previously^132^. Larvae were euthanized by rapid cooling on a mixture of ice and water between 2-4°C for a minimum of 30 minutes after complete cessation of movement was observed. Genotyping was performed on all individual fish at the conclusion of each sleep assay. Either restriction digest or headloop PCR methods were used to validate mutations^37,133^. Primers, restriction enzymes, and PCR parameters for genotyping are listed in **Supplementary Table 1**. All primers were run on a 2% agarose gel and sequence verified using Sanger sequencing to verify the target region.

### Data collection and analysis for sleep phenotyping in zebrafish

All embryos and larvae were housed in an incubator at 28.5°C, with lights on at 9am (ZT0) and lights off at 11pm (ZT14) prior to data collection. Dead embryos and chorion were removed daily until 5 dpf. At 5 dpf, crispants and scrambled-injected sibling controls were screened for gross morphological deficits and swim bladder inflation. Only healthy larvae were pipetted into individual wells of a 96-well plate and placed into a ZebraBox (ViewPoint Life Sciences) for automated video monitoring. Sleep-wake behaviors were measured through automated video-tracking as described previously^37,132,134^. Two biological replicates were run for *dpyda/b*, *myrf*, and *fbxw7* using different clutches (sibling-matched within clutch) of embryos and well placement was flipped for each experiment to minimize location bias across experiments. One clutch was run for *dpys* and *upb1*. Data were processed using custom R scripts (https://github.com/amber-zim/Zebrafish-sleep-analysis-from-Viewpoint-2026) developed from MATLAB code generated previously^134^ to calculate the sleep and activity parameters. For *dpyda/b*, *myrf*, and *fbxw7*, two independent experiments were pooled. Behavioral fingerprints (z-score deviations from controls) were calculated using ZOLTAR^62^ (https://francoiskroll.shinyapps.io/zoltar/) and are available within the app. Data were plotted for each clutch separately in **Extended Data** to show consistency and as the average of the two clutches in **Figure 3f-g**. ZOLTAR uses a threshold of 0.1 second of movement per min to define sleep. Thus, to align our data with the previous drug screen data analyzed using ZOLTAR, we used this same threshold. However, for all other analyses, a threshold of 0.5 sec/min is used to define sleep as this threshold was optimized for our system and is aligned with the threshold previously reported to differentiate true sleep from quiet wake^134^. This had a negligible effect on our results, and we found 0.5 to be more restrictive to detecting significant sleep phenotypes. All individual day data are provided in **Extended Data**. Data averaged across both days are provided in the main figures. For averaged day data, normality was tested using the D’Agostino-Pearson omnibus test in Graphpad Prism v10. Normal data were analyzed by unpaired student’s t-test and non-parametric data were analyzed by Mann Whitney test unless stated otherwise. For individual day data, Wilcoxon rank sum test was used followed by FDR (Benjamini-Hochberg) multiple comparisons correction across all measures. In the cases of *fbxw7* and *upb1* crispants where a linear regression model was used to control for effects on waking activity; the following model was implemented in R:

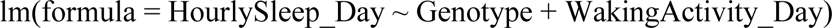

### Drug administration in zebrafish

Clemastine fumarate was purchased from Santa Cruz Biotech. For clemastine experiments, dose selection was performed on wild-type fish. Six doses of clemastine were prepared by serial dilution from 20 mM to 0.625 mM [20, 10, 5, 2.5, 1.25, 0.625 mM] from a stock solution dissolved in 100% DMSO as previously described^63,135^. Zebrafish larvae were pipetted into individual wells of a 96-well plate at 3 dpf after naturally shedding their chorion to not interfere with normal development. 650 µL of zebrafish embryo media (E3) was used to fill each well flush with the top; then drugs were applied to generate a 1:1000 dilution into the media [e.g. 0.65 µL of 1.25 mM clemastine solution pipetted into 650 µL for a final concentration of 1.25 µM in 0.1% DMSO]. 100 nM melatonin (Sigma-Aldrich) dissolved in DMSO was used as a positive control as it has been shown to potently induce sleep in zebrafish^135^. DMSO (0.1%) was applied to control wells as it was used to dissolve clemastine and can increase swimming activity at concentrations >0.5%^136^. Drugs were applied at 3, 4, and 5 dpf. E3 was filled to the top of each well and the drug was added to the top. Approximately one-half to one-third of the media evaporates each day and is topped up with fresh media each day. To minimize disruption to fish, the remaining media was not removed; therefore, concentrations may have accumulated across multiple days, which is why we used lower doses than previously described^63^. Sealing plates generates bubbles and obscures video tracking and may cause drug leakage across wells^134^; thus, this method proved more reliable in our experimental setting. No drug was applied from days 6-8 post fertilization while behavior was being tracked; however, residual drug was likely maintained in the wells following media changes.

For the melatonin-only experiment, 1 mM melatonin stock solution was made by dissolving melatonin in 100% DMSO. This solution was diluted 1:1000 by pipetting 0.65 µL into individual wells containing 650 µL of E3 media for a final concentration of 1 µM melatonin in 0.1% DMS0. Melatonin or DMSO was administered at 6 and 7 dpf at ZT2. Sleep and activity were examined in the 1-hour post administration relative to the 1-hour pre administration.

### Whole mount *in situ* hybridization chain reaction (HCR)

HCR experiments were performed in *Tg(elavl3:H2B-GCaMP6s)jf5Tg* larvae homozygous for the *mitfa^w^*^2^ allele (nacre). Larvae were fixed in 4% PFA at 6 dpf for approximately 24 hours then washed with 1× PBS 3 times for 5 mins. Whole larvae were washed for 5 minutes 3 times with 1× PBST. *In situ* hybridization protocol was performed as previously described^137,138^. Custom probes were designed using initiator sequences previously described^139^ to generate 15 probes pairs for *dpyda*, *ptprz1b,* and *sox2*. Probe design was performed using publicly available code from github: https://github.com/rwnull/insitu_probe_generator. HCR buffers and B amplifiers (B1-Alexa Fluor 594; B3-Alexa Fluor 546; B5-Alexa Fluor 647) were purchased from Molecular Instruments (https://www.molecularinstruments.com/).

### Confocal imaging of zebrafish

Whole zebrafish larvae were mounted in 1% low melting agarose against Corning glass coverslips and imaged using a Zeiss LSM 980 Airyscan 2 confocal microscope equipped with either a ZEISS C-Aprochromat 10×/0.45 W M27 objective or a LD LCI Plan-Apochromat 25x/0.8 Imm Corr DIC M27 objective. Laser settings were held consistent for every image. The laser wavelength was 639 nm for B5-AF 647; 561 nm for B3-Alexa Fluor 546; 594nm for B1-AF594; 488 nm for GCaMP6s. Z-stacks were taken at 1µm intervals. Confocal z-stacks were visualized using FIJI. Maximum intensity projections were created for each image used to examine whole brain expression. Orthogonal projections were generated using FIJI.

### Fly stocks and maintenance

The UAS-*su(r)-*RNAi HMC03568 line was obtained from the Bloomington *Drosophila* Stock Center (BL 53339). *nSyb*-GAL4, *repo*-GAL4 were gifts from L. Griffith, Brandeis, 9-137-GAL4 was a gift from R. Bainton, UCSF, NP2222-GAL4 and MZ0709-GAL4 (experimental stock used for this cross was GAL80ts; MZ0709-GAL4) were a gift from M. Freeman, OHSU.

All fly stocks were maintained on a molasses-based food source at room temperature. For experimental crosses, flies were kept on molasses-based food, except during the sleep measurements (see details below). Crosses were maintained on a 12-h:12-h light/dark cycle at 25°C. For each experiment, 3-5 virgin females were selected for and crossed with 3-5 males.

### Sleep analysis in *Drosophila*

For sleep experiments, males and females were selected for at 3-5 days post eclosion. Individual flies were placed into glass tubes containing 5% sucrose and 2% agarose. Sleep and locomotor activity data were collected in 1-minute bins for a minimum of 3 days using multibeam *Drosophila* activity monitors (Trikinetics DAM5H), which have 15 infrared beams that provide high precision activity monitoring. Following a 24-hour acclimation period, activity across 48 hours was included in analyses and sleep-wake data were averaged across the two days. Sleep in *Drosophila* is defined as 5 continuous minutes of inactivity. Flies were examined upon removal from the DAM, and any fly that was dead was not included in the analysis. Each experimental group was compared against two controls (+/UAS *su(r)* RNAi and GAL4/+ only) to control for background-specific effects using one-way ANOVA and Holm-Sidak multiple comparisons test. To simplify data presentation, sleep duration is plotted as the difference between experimental and control groups as described previously^77^. To visualize all activity and sleep measures, effect sizes (Cohen’s *d*) were calculated as experimental group relative to each control.

### Statistical analysis of sleep across species

To standardize behavioral data across species, we calculated the effect sizes as standardized mean difference (Cohen’s *d*) for each genetic mutant versus its assay control. For consistency with other species, zebrafish data were pooled from two independent clutches and Cohen’s *d* was calculated for all *dpyda/b* crispants versus all controls. For *Drosophila*, *su(r)* RNAi lines were compared against each parental control (+/UAS *su(r)* RNAi and GAL4/+) separately. Effect sizes for mouse were taken from Table 1 in *Keenan et al. 2021*^8^ in which sleep was measured by EEG. For normal data, Cohen’s *d* and z-scores are equivalent and can be compared to the behavioral fingerprint data. Absolute values of these differences equal to 0.2, 0.5, or >0.8 can be interpreted as small, medium and large effects, respectively^140^.

### Lipid droplet imaging in *Drosophila*

Male and female flies were collected at 5-10 days post eclosion, decapitated, and brains were dissected on ice in 1× Phosphate Buffer Saline (PBS). Afterward, they were fixed in 4% PFA (diluted in 1× PBS), washed, and whole brains were stained using 1µg/ml BODIPY^493/503^ (diluted in 1× PBST, BODIPY^493/503^, Fisher-D3922) and mounted in Vectashield Antifade Mounting Media (Vector Laboratories, H-1400) on glass slides with coverslips. Whole brains were imaged on Leica SP5 confocal microscope, using tiled z-stacks under a 40× objective. LDs were quantified in ImageJ as previously described^77^. For each z-slice the background was subtracted using a convolution background filter before a mask was applied capturing the area of the brain. Within that area, LD total counts and size were captured (particle size limits of 2-150). Count and normalized LD size were compared between *repo*-GAL4>UAS-*su(r)* RNAi and its two controls (+/UAS *su(r)* RNAi and GAL4/+ only) to control for background-specific effects using one-way ANOVA and Holm-Sidak multiple comparisons test.

### Public data curation and analysis

GWAS sentinel variants were collected from Sleep Disorders Knowledge Portal (https://sleep.hugeamp.org/) for sleep duration^2^. Linkage disequilibrium *r*^2^ values were computed using LDLink^88^ (https://ldlink.nih.gov/) with the LDProxy tool against GRCh37 (hg19) with the ancestry set to CEU and a calculation window of 500Kb and sorted by FORGEdb^89^ score. eQTLGen blood cis-eQTL data was obtained from https://forgedb.cancer.gov/.

Single cell RNA sequencing data from mouse were obtained from *Tabula Muris*^19^ and Bandler et al.^23^. Data from humans were collected from Velmeshev et al.^22^. Data from zebrafish were collected from Daniocell^43^. Data from *Drosophila* were collected from Fly Cell Atlas^78^.

Cis-Regulatory atlases were obtained from: *Drosophila*^95^: visualized using the UCSC Genome Browser track hub: http://ucsctracks.aertslab.org/papers/FlyBrain/hub.txt on dm6 coordinates. Zebrafish^94^: visualized using WashU EpiGenome Browser on danrer10 coordinates. Mouse^93^: visualized at https://catlas.org/catlas_mousebrain/#!/igv on mm10 coordinates Human^87^: visualized using UCSC Genome browser on hg19 coordinates. Genome builds were chosen based on available cis-regulatory data.

## Acknowledgements

Authors would like to thank the CHOP Zebrafish Core (RRID:SCR_028414) for assistance in zebrafish care, maintenance, and CRISPR experimentation and the Institute of Structural Biology and Beckman Center for Cryo-EM at the University of Pennsylvania (RRID: SCR_022375). Authors also thank Aalim Weljie and Dania Malik for their assistance in LC-MS experimentation and analysis and Miyoko Imamura and Jie Lian for their assistance with animal husbandry and phenotyping.

## Funding

The work was supported by NIH grant T32 HL170968, R01 HL178074, P01 HL094307, and P01 HL160471. S.D. is supported by funding from the IRACDA program at the University of Pennsylvania from NIGMS (K12GM081259). F.K. is supported by the Fondation pour la Recherche Médicale grant SPF202309017505. S.F.G is supported by NIH awards R01 AG057516 and R01 HD056465 and the Daniel B. Burke Endowed Chair. J.R. is supported by a Wellcome Trust Investigator Award (#217150/Z/19/Z), a BBSRC grant (BB/X01536X/1) and a European Research Council Advanced Grant. A.I.P. is The John L. Miclot Professor of Medicine at the University of Pennsylvania; funds for this endowment were provided by the Philips Respironics Foundation.

## Author Contributions

Conceptualization: A.J.Z, S.D., A.S., J.R. A.I.P.

Methodology: A.J.Z., S.D. E.A., F.K., F.D.B., J.D., B.T.K., X.G.

Investigation: A.J.Z., S.D., E.A., T.R.T. Z.Y.S., F.D.B., J.D.

Visualization: A.J.Z.

Supervision: S.F.A.G., A.S., J.R., A.I.P.

Writing—original draft: A.J.Z, A.I.P

Writing—review & editing: All authors

## Declaration of competing interests

Authors have nothing to disclose.

## Data availability

Data are available from the corresponding author upon reasonable request.

## Code availability

The code to analyze zebrafish sleep and activity has been made available at https://github.com/amber-zim/Zebrafish-sleep-analysis-from-Viewpoint-2026.

## Notes

### Competing Interest Statement

The authors have declared no competing interest.

