## Extended Data for "*DPYD* links glial lipids and myelination to sleep duration"

Zimmerman et al.

Extended Data Figures 1-12

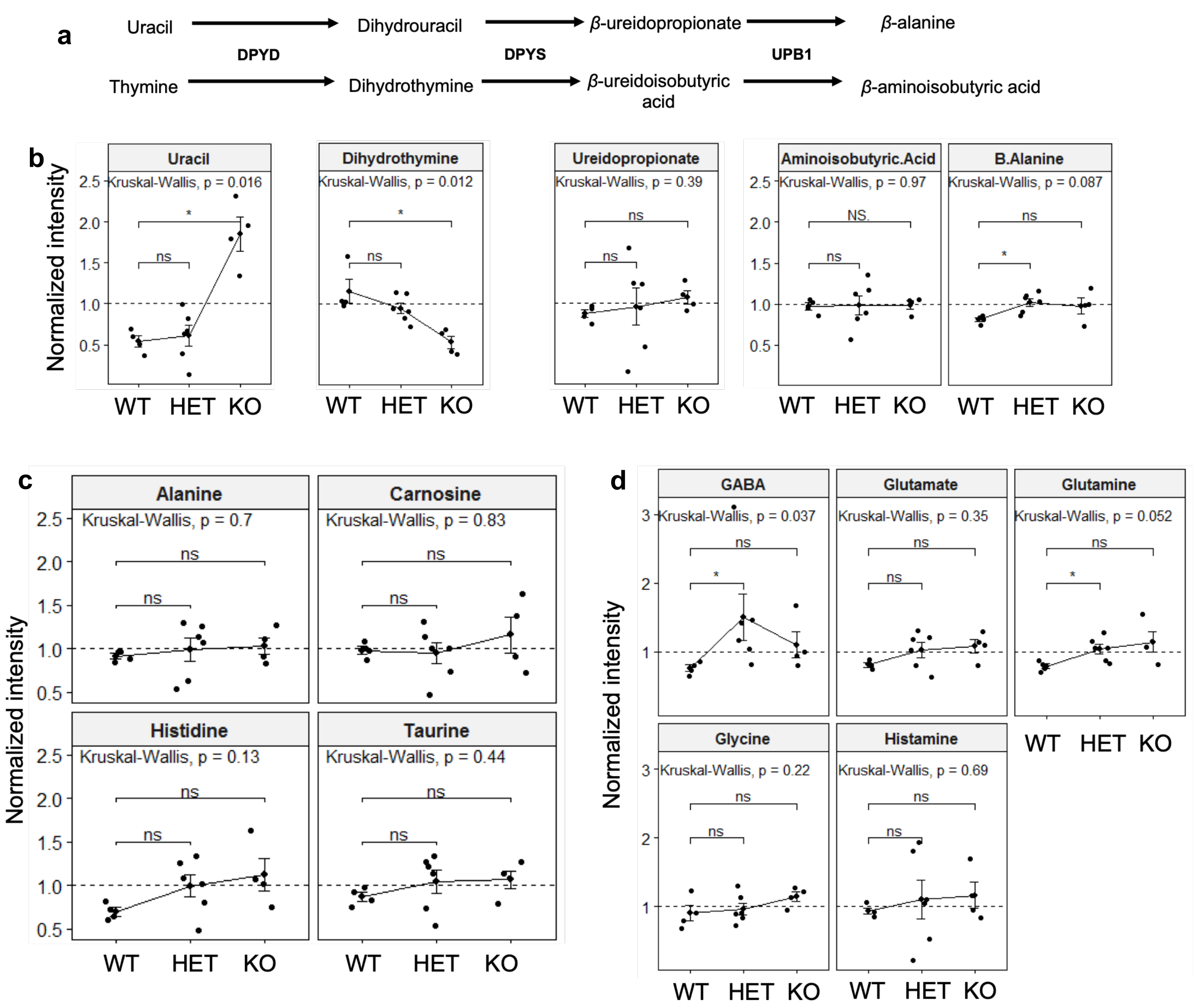


**Extended Data Fig. 1. Brain metabolites assayed by liquid chromatography mass spectrometry. a.** Pyrimidine catabolism pathway showing the enzymes and respective intermediates generated at each step. **b.** Metabolites tested in the pyrimidine catabolic pathway from piriform cortex samples taken from *Dpyd* homozygous knockouts (KO), heterozygous (HET), and wild-type (WT) mice. Additional metabolites related to $\beta$-alanine (**c**) and neurotransmission (**d**). Thymine, dihydrouracil, and $\beta$-ureidoisobutyric acid were not detected in these samples. Data points represent individual animals overlaid with group mean values connected across genotypes, and error bars denote the standard error of the mean (Mean ± SEM). Statistical significance for Mann-Whitney U post-hoc comparisons is indicated as follows: *$P < 0.05$ and ns (not significant, $P \geq 0.05$). Overall Kruskal-Wallis $P$-values along with post-hoc significance markers are annotated on each panel.

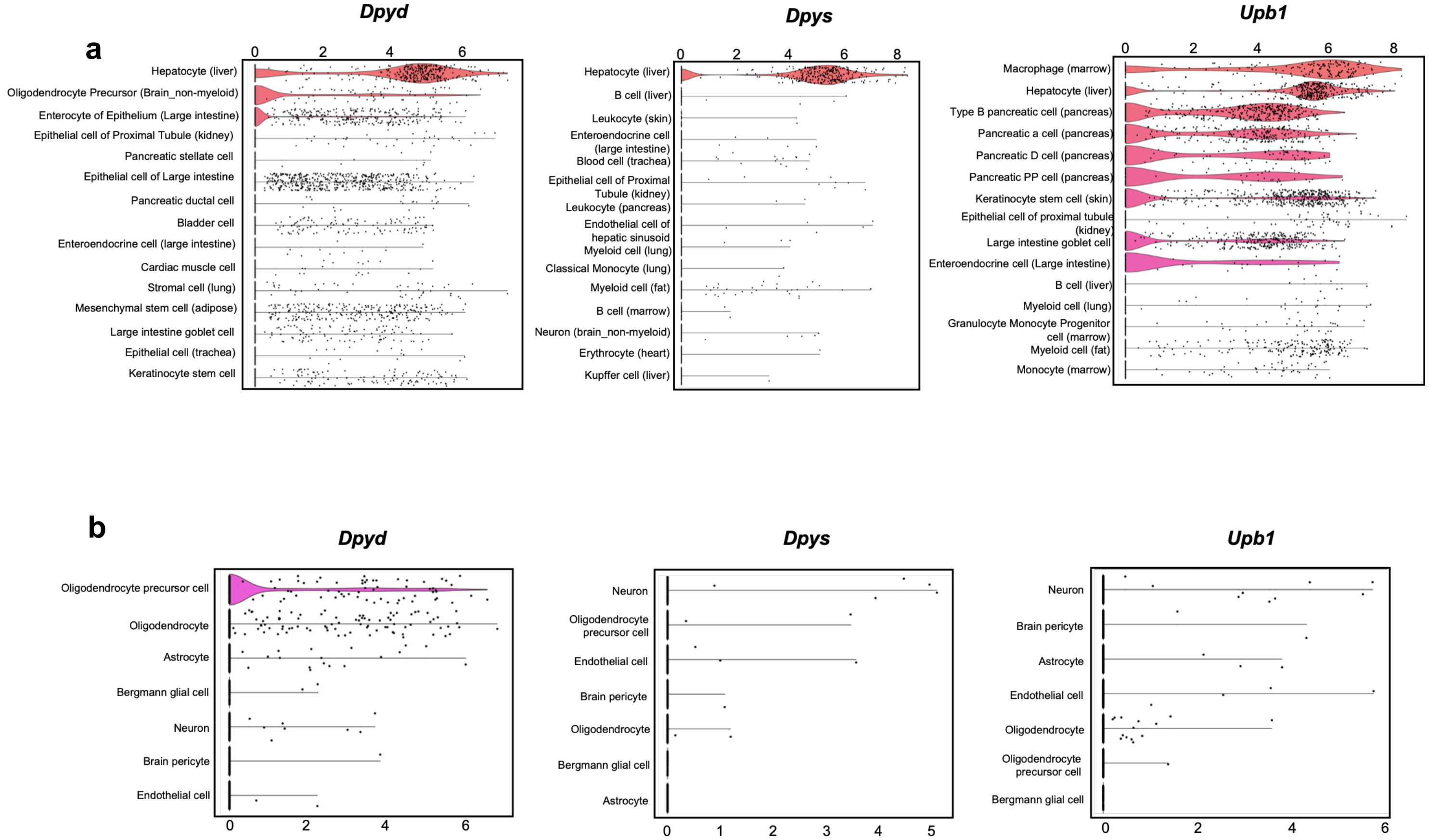


**Extended Data Fig 2. Single cell RNA expression of pyrimidine catabolic enzymes in mouse tissue**. RNA-sequencing data taken from *Tabula Muris* (https://tabula-muris.sf.czbiohub.org). **a.** Top 15 tissues ranked by highest (top) to lowest (bottom) gene expression across all tissues. **b.** Brain (non-myeloid tissue) specifically showing expression ranked from highest (top) to lowest (bottom). Expression values presented ln(1+CPM).

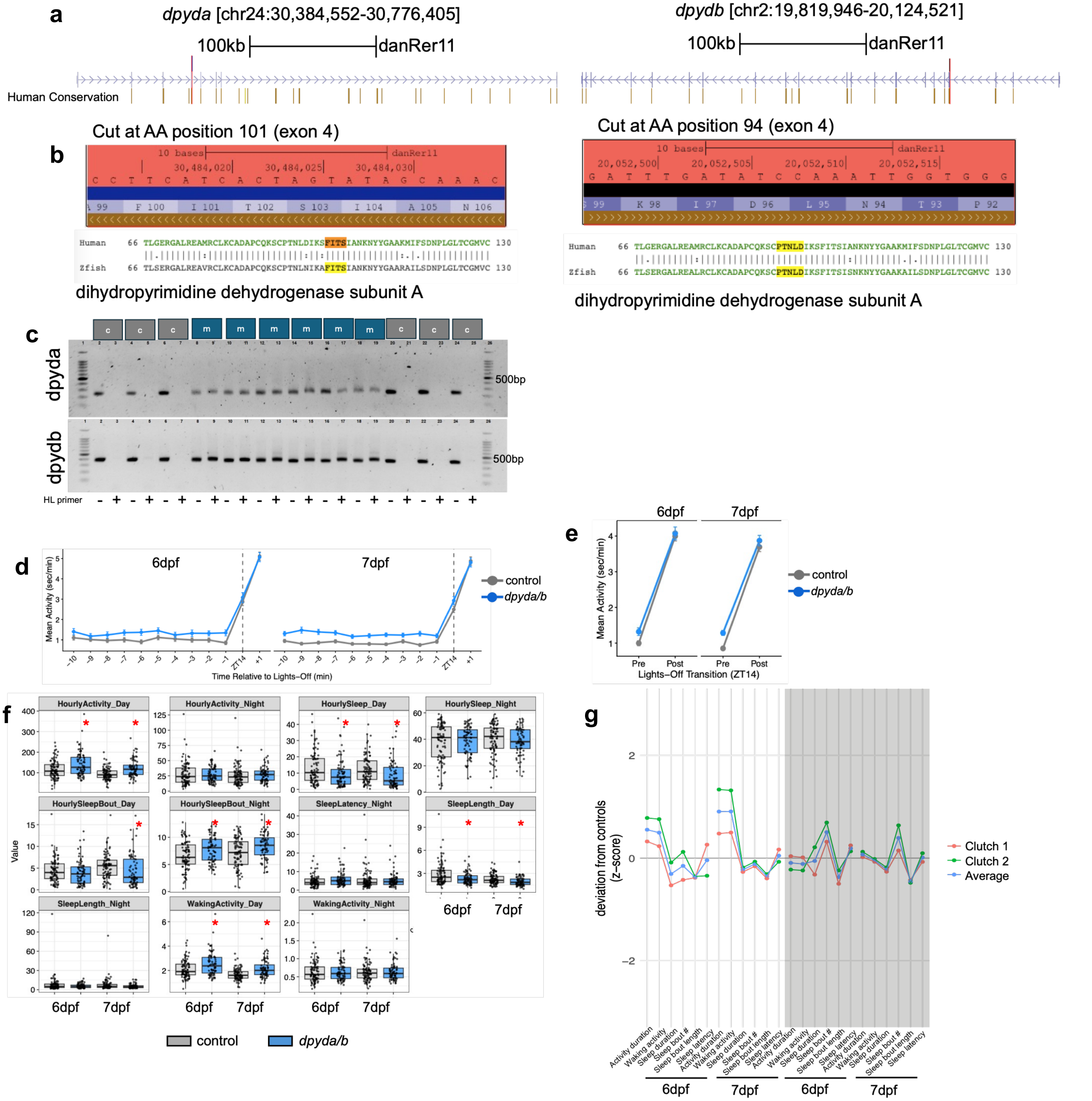


**Extended Data Fig. 3. Mutation of *dpyda/b* in zebrafish. a.** UCSC browser image of the *dpyda* and *dpydb* genes showing the location of the guideRNA target (red line). Human conservation blocks are colored by corresponding human chromosome. **b.** Zoomed in region of the UCSC browser to the red highlight indicating the guideRNA along with the amino acid (AA) sequence and the exon targeted. The AA conservation between human and zebrafish (zfish) is shown from DIOPT. Straight bars indicate conservation. The protein domain (listed by DIOPT) targeted is labeled below the AA conservation. **c.** Genotype validation using headloop PCR as described in **Methods**. C = control, m = mutant. HL primer + indicates addition of the headloop primer; HL- indicates only standard forward and reverse primers. If wild-type DNA is present, HL primer will suppress amplification. If sequence is mutated, HL primer will not bind and PCR amplicon will be present. **d.** Dark transition arousal response (mean and SEM) on days 6 and 7 post fertilization. Data plotted as time relative to lights off (ZT14). **e.** Quantification (mean and SEM) of arousal response in d. Data represent the change in activity in the 2 minutes after ZT14 relative to the average of the 10 minutes before ZT14. **f.** Behavioral measures plotted separately by day. Data plotted as box plots representing median and quartiles with individual fish plotted. (n = 72 *dpyda/b* crispants and 88 sibling negative controls) data are pooled from two independent experiments. Red asterisks indicate significance following FDR correction of the Wilcoxon rank sum test. **g.** Behavioral fingerprints from two independent clutches. Data are presented as z-score deviation relative to sibling and plate-matched controls for each separate assay (clutch). The average of the combined rounds is used in main Figures 2 and 3.

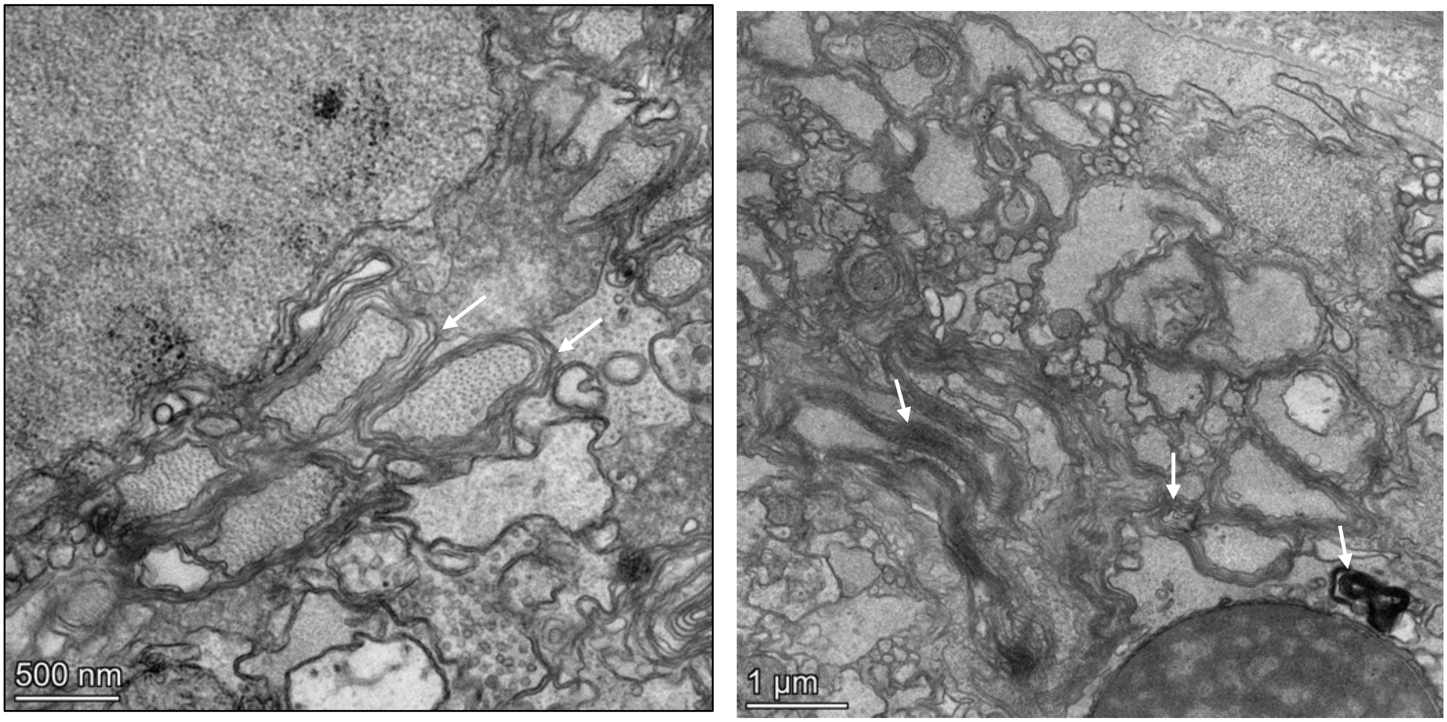


**Extended Data Fig. 4.** Representative electron micrographs of decompacted (left) and aberrant (right) myelin sheaths in the spinal cord of 8 dpf *dpyda/b* crispant zebrafish. White arrows point to specific examples.

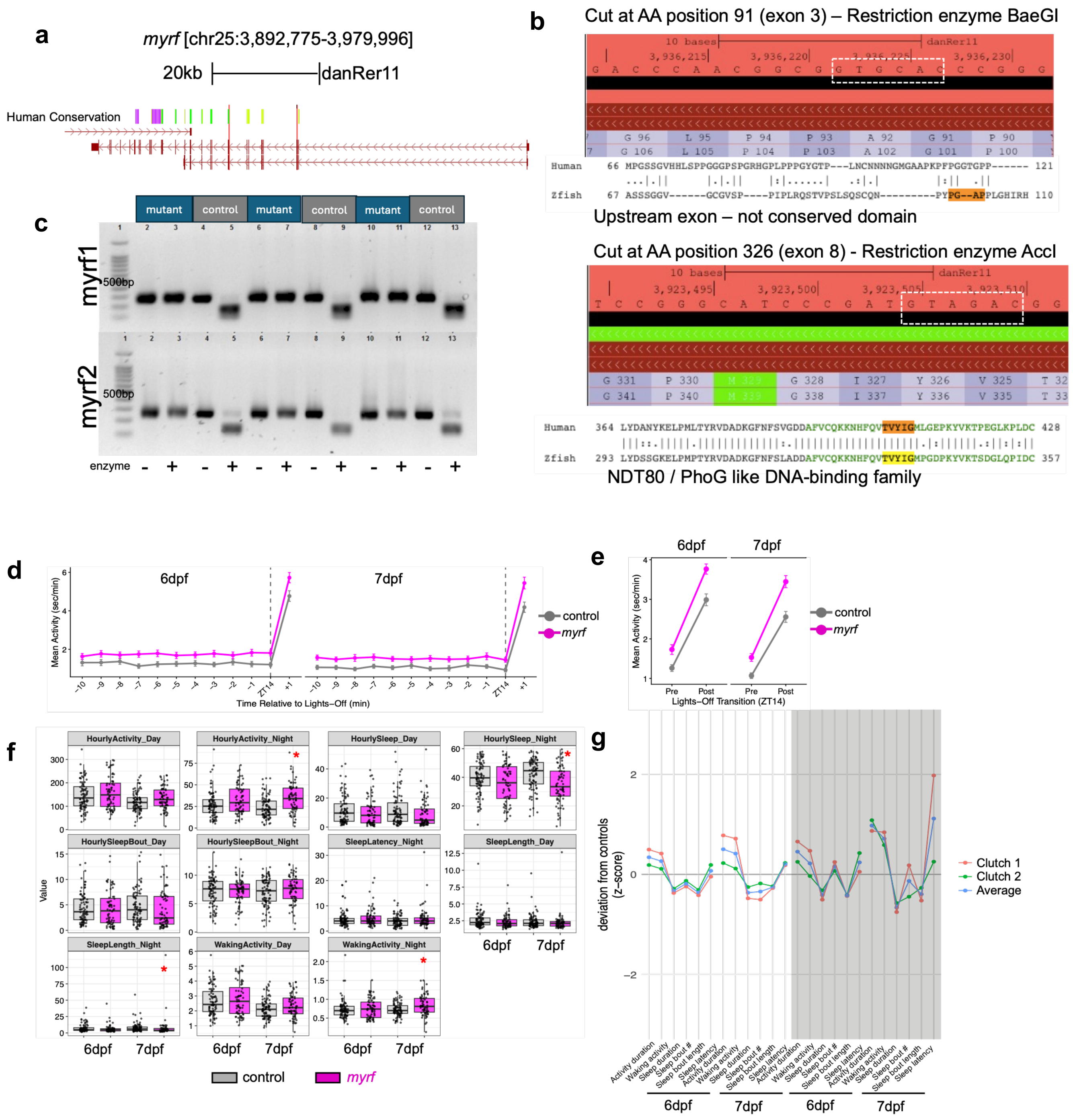


**Extended Data Fig. 5. Mutation of *myrf* in zebrafish. a.** UCSC browser image of the *myrf* genomic region showing the location of the guideRNA targets (red lines). Human conservation blocks are colored by corresponding human chromosome. **b.** Zoomed in region of the UCSC browser to the red highlight indicating the guideRNA along with the amino acid (AA) sequence and the exon targeted. The restriction enzyme cut site is outlined by a white dashed box. The AA conservation between human and zebrafish (zfish) is shown from DIOPT. Straight bars indicate conservation. The protein domain (listed by DIOPT) targeted is labeled below the AA conservation. **c.** Genotype validation using restriction digest PCR as described in methods. If wild-type DNA is present, PCR product will be digested. If sequence is mutated, PCR product will not be digested. **d.** Dark transition arousal response (mean and SEM) on days 6 and 7 post fertilization. Data plotted as time relative to lights off (ZT14). **e.** Quantification (mean and SEM) of arousal response in d. Data represent the change in activity in the 2 minutes after ZT14 relative to the average of the 10 minutes before ZT14. **f.** Behavioral measures plotted separately by day. Data plotted as box plots representing median and quartiles with individual fish plotted. (n = 68 *myrf* crispants and 86 sibling negative controls) data are pooled from two independent experiments. Red asterisks indicate significance following FDR correction of the Wilcoxon rank sum test. **g.** Behavioral fingerprints from two independent clutches. Data are presented as z-score deviation relative to sibling and plate-matched controls for each separate assay (clutch). The average of the combined rounds is used in main Figures 3.

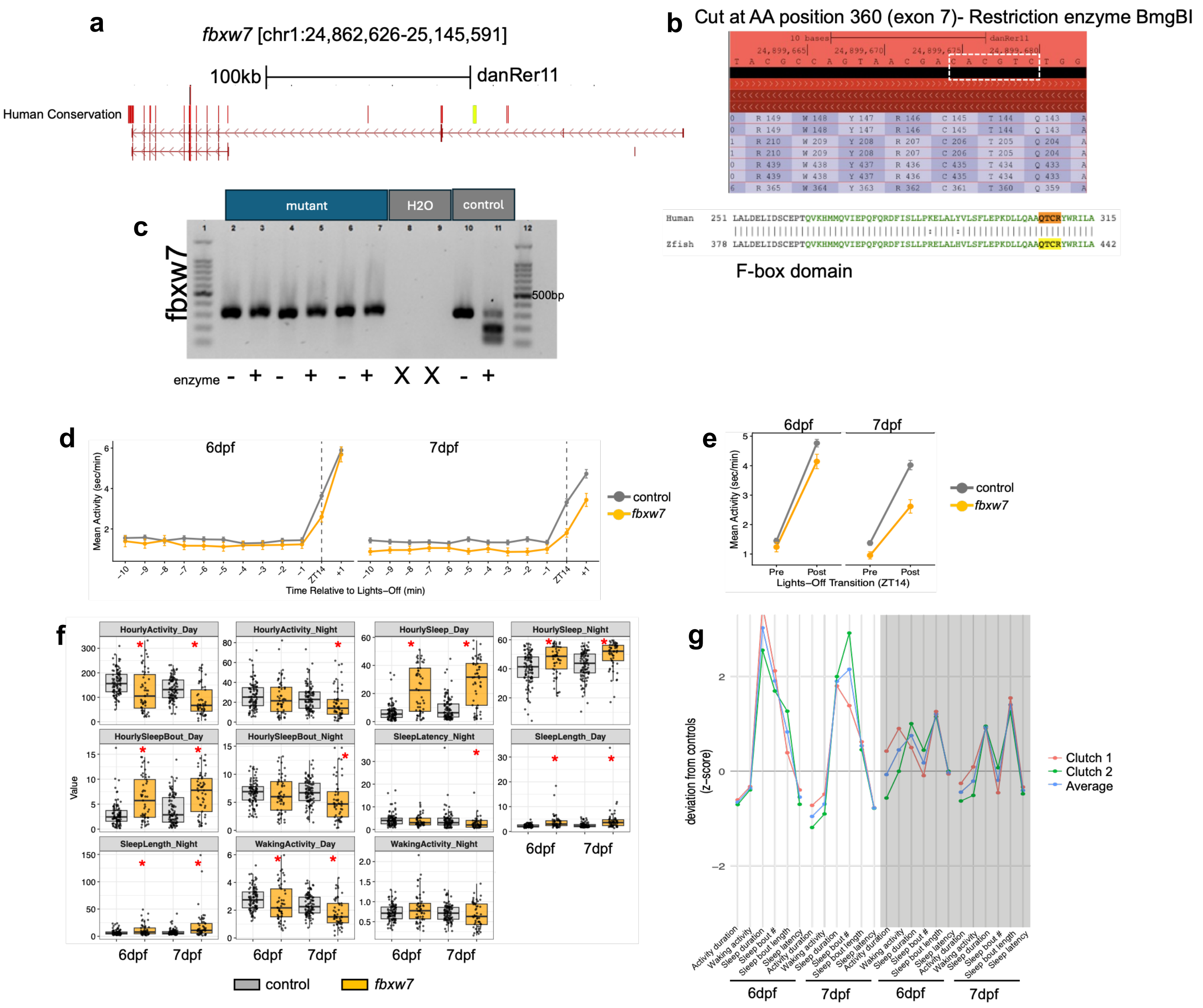


**Extended Data Fig. 6. Mutation of *fbxw7* in zebrafish**. **a.** UCSC browser image of the *fbxw7* gene showing the location of the guideRNA (red line). Human conservation blocks are colored by corresponding human chromosome. **b.** Zoomed in region of the UCSC browser to the red highlight indicating the guideRNA along with the amino acid (AA) sequence and the exon targeted. The restriction enzyme cut site is outlined by a white dashed box. The AA conservation between human and zebrafish (zfish) is shown from DIOPT. Straight bars indicate conservation. The protein domain (listed by DIOPT) targeted is labeled below the AA conservation. **c.** Genotype validation using restriction digest PCR as described in methods. If wild-type DNA is present, PCR product will be digested. If sequence is mutated, PCR product will not be digested. **d.** Dark transition arousal response (mean and SEM) on days 6 and 7 post fertilization. Data plotted as time relative to lights off (ZT14). **e.** Quantification (mean and SEM) of arousal response in d. Data represent the change in activity in the 2 minutes after ZT14 relative to the average of the 10 minutes before ZT14. **f.** Behavioral measures plotted separately by day. Data plotted as box plots representing median and quartiles with individual fish plotted. (n = 53 *fbxw7* crispants and 95 sibling negative controls) data are pooled from two independent experiments. Red asterisks indicate significance following FDR correction of the Wilcoxon rank sum test. **g.** Behavioral fingerprints from two independent clutches. Data are presented as z-score deviation relative to sibling and plate-matched controls for each separate assay (clutch). The average of the combined rounds is used in main Figures 3.

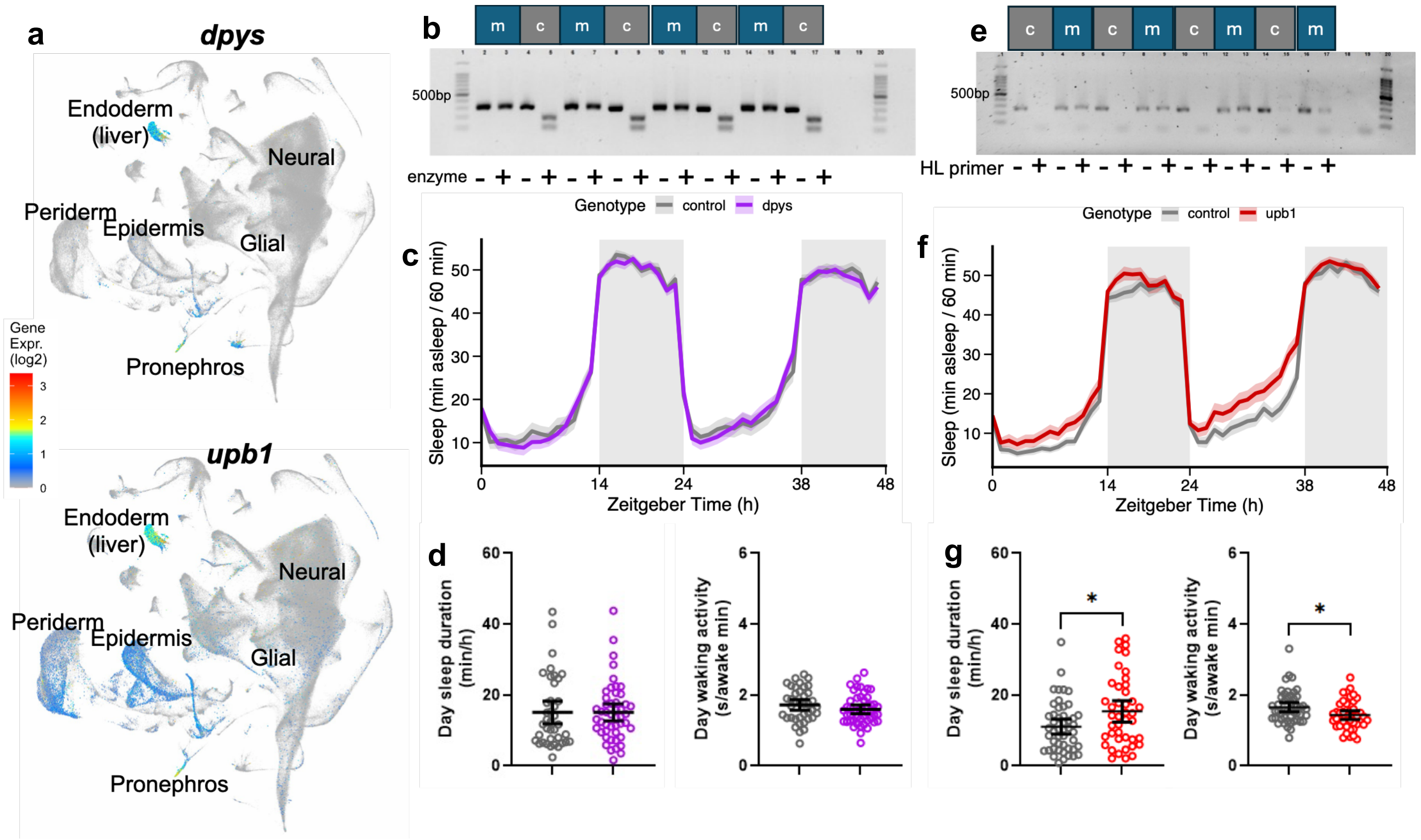


**Extended Data Fig. 7. Mutation of zebrafish genes encoding liver-expressed pyrimidine catabolic enzymes *dpys* and *upb1*. a.** Single-cell RNA-sequencing data from Daniocell showing *dpys* and *upb1* expression in all cell types. **b.** *dpys* genotype validation. **c.** Sleep trace representing mean and SEM for *dpys* crispants (n = 48) and negative controls (n = 39). **d.** Quantification of daytime sleep duration and waking activity in *dpys* crispants and negative controls. **e.** *upb1* genotype validation. **f.** Sleep trace representing mean and SEM for *upb1* crispants (n = 48) and negative controls (n = 42). **g.** Quantification of daytime sleep duration and waking activity in *upb1* crispants and negative controls. Quantification plots show mean and 95% CI. **P*<0.05 determined by non-parametric Mann Whitney test. In b and e: Blots show PCR amplification of mutants (m) and controls (c) by restriction digest (enzyme) or headloop (HL) PCR.

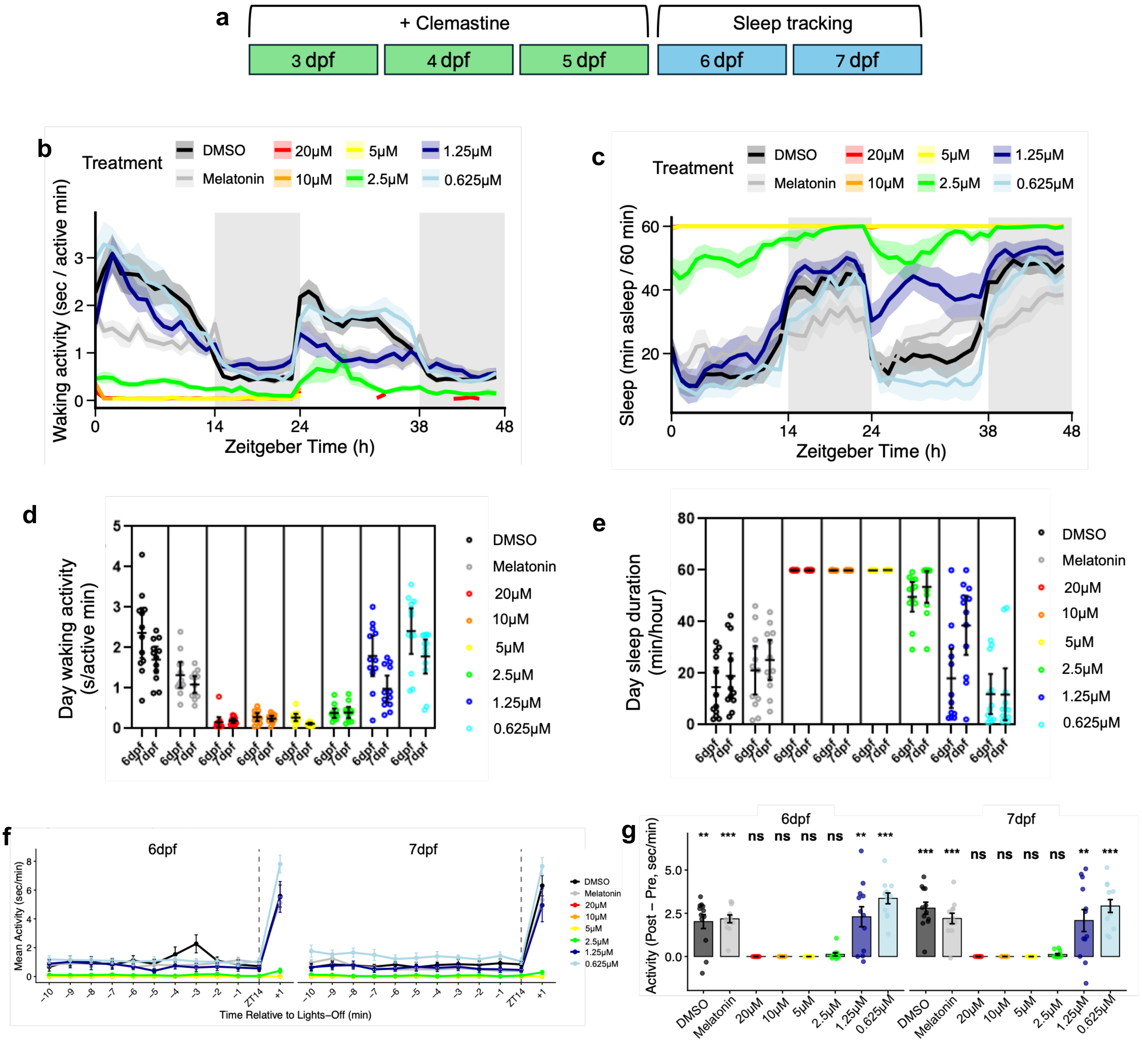


**Extended Data Fig. 8. Clemastine dose-response assessment in wild-type zebrafish. a.** Clemastine drug administration paradigm. Mean and SEM waking activity (**b**) and sleep (**c**) in wild-type zebrafish larvae collected from days 6 and 7 post fertilization. Quantification (mean and 95% CI) of day waking activity (**d**) and sleep duration (**e**) on days 6 and 7 post fertilization. n = 12 per treatment. **f.** Dark transition (mean and SEM) arousal response on days 6 and 7 post fertilization. Data plotted as time relative to lights off (ZT14). **g.** Quantification (mean and SEM) of the arousal response in d. Data represent the change in activity in the 2 minutes after ZT14 relative to the average of the 10 minutes before ZT14. **P*<0.05, ***P*<0.01, ****P*<0.001, ns = not significant, determined by paired t-test comparing pre vs. post followed by FDR correction using the Benjamini-Hochberg method.

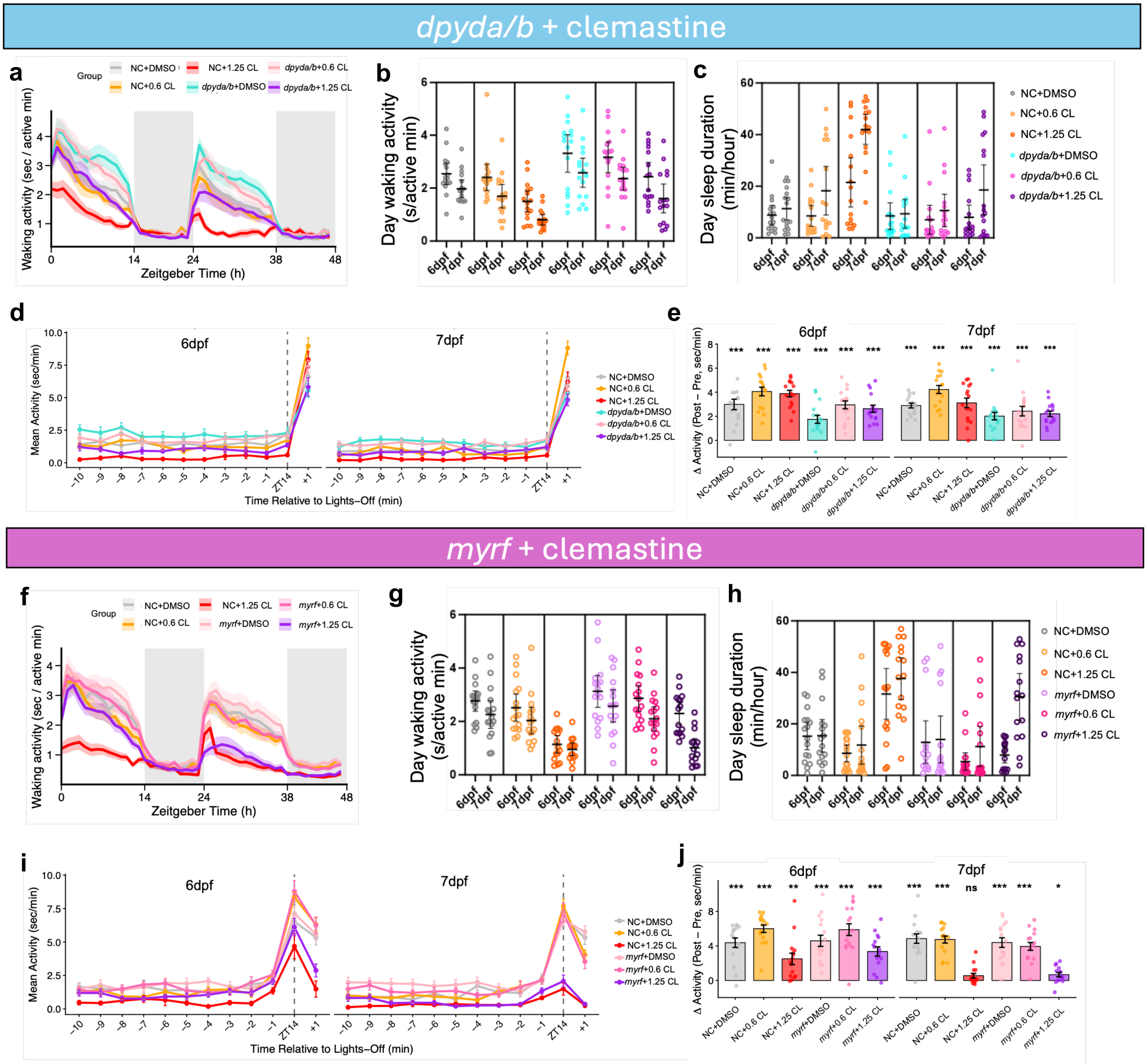


**Extended Data Fig. 9. Clemastine administration in *dpyda/b* and *myrf* crispants. a.** Mean and SEM waking activity in *dpyda/b* crispant and sibling negative controls (n = 16/ group). Data collected from days 6 and 7 post fertilization. Genotype and clemastine dose are indicated. CL = clemastine. Quantification (mean and 95% CI) of day waking activity (**b**) and sleep duration (**c**). **d.** Dark transition arousal response (mean and SEM) on days 6 and 7 post fertilization. Data plotted as time relative to lights off (ZT14). **e.** Quantification (mean and SEM) of the arousal response in d. **f.** Mean and SEM waking activity in *myrf* crispant and sibling negative controls. Quantification (mean and 95% CI) of day waking activity (**g**) and sleep duration (**h**). **i.** Dark transition arousal response (mean and SEM) for *myrf* crispants and negative controls given drug (n = 15-16/group). **j.** Quantification (mean and SEM) of the arousal response in i. For e and j, data represent the change in activity in the 2 minutes after ZT14 relative to the average of the 10 minutes before ZT14. **P*<0.05, ***P*<0.01, ****P*<0.001, ns = not significant, determined by paired t-test comparing pre vs. post followed by FDR correction using the Benjamini-Hochberg method.

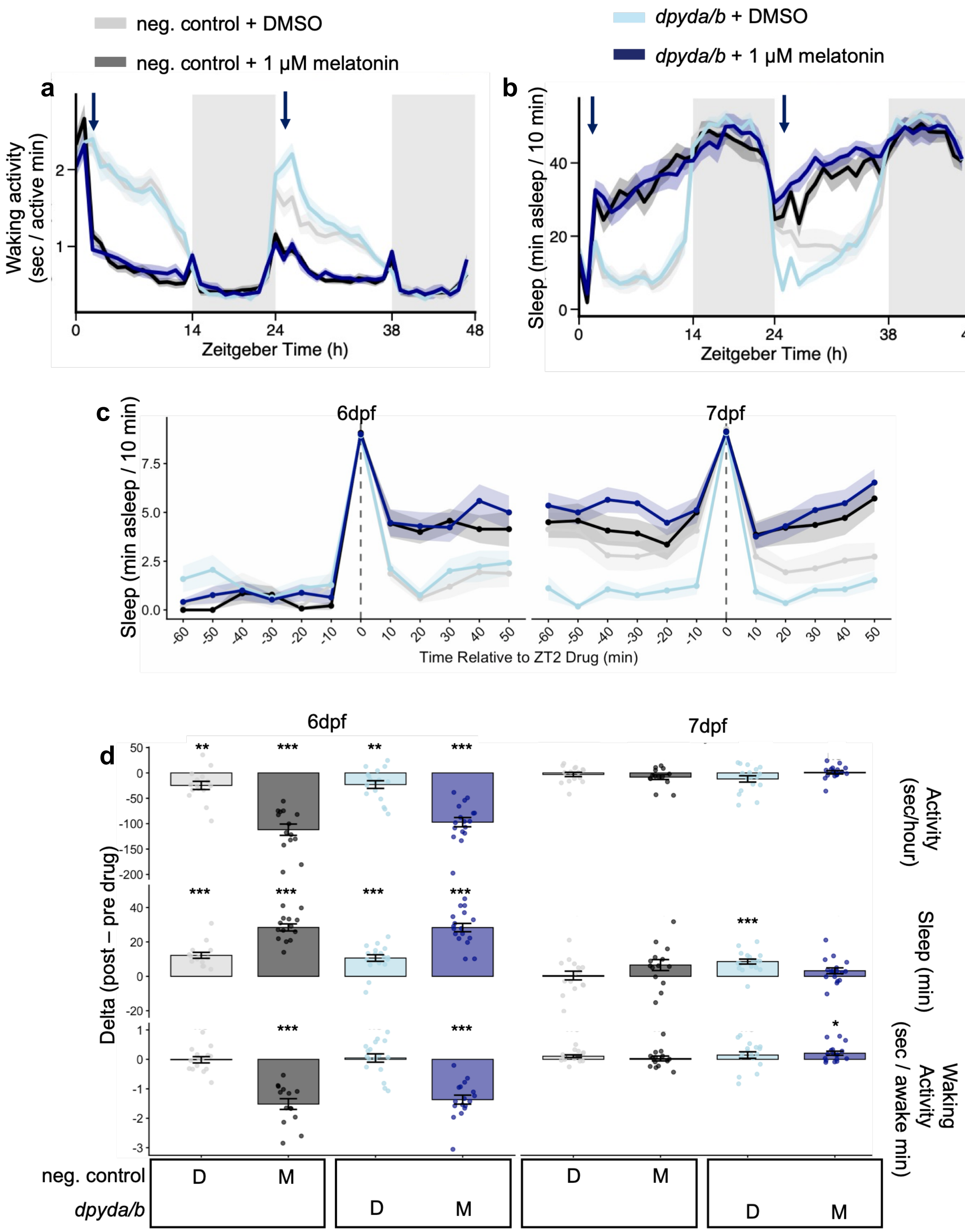


**Extended Data Fig. 10. Sleep and activity parameters in *dpyda/b* crispants following melatonin administration.** Mean and SEM waking activity (**a**) and sleep duration (**b**) in *dpyda/b* crispants and sibling negative controls given DMSO or 1 μM melatonin at ZT 2 (blue arrows). **c.** Behavioral response (mean and SEM) to drug administration in the 60 minutes pre and 60 minutes post administration. **d.** Quantification (mean and SEM) of the behavioral response (Delta of the 60 minutes post relative to 60 minutes pre drug administration). n = 15 control + DMSO, 14 control + melatonin, 17 *dpyda/b* + DMSO, 17 *dpyda/b* + melatonin. **P*<0.05, ***P*<0.01, ****P*<0.001, determined by paired t-test comparing pre vs. post followed by FDR correction using the Benjamini-Hochberg method. In d, D = DMSO, M = melatonin.

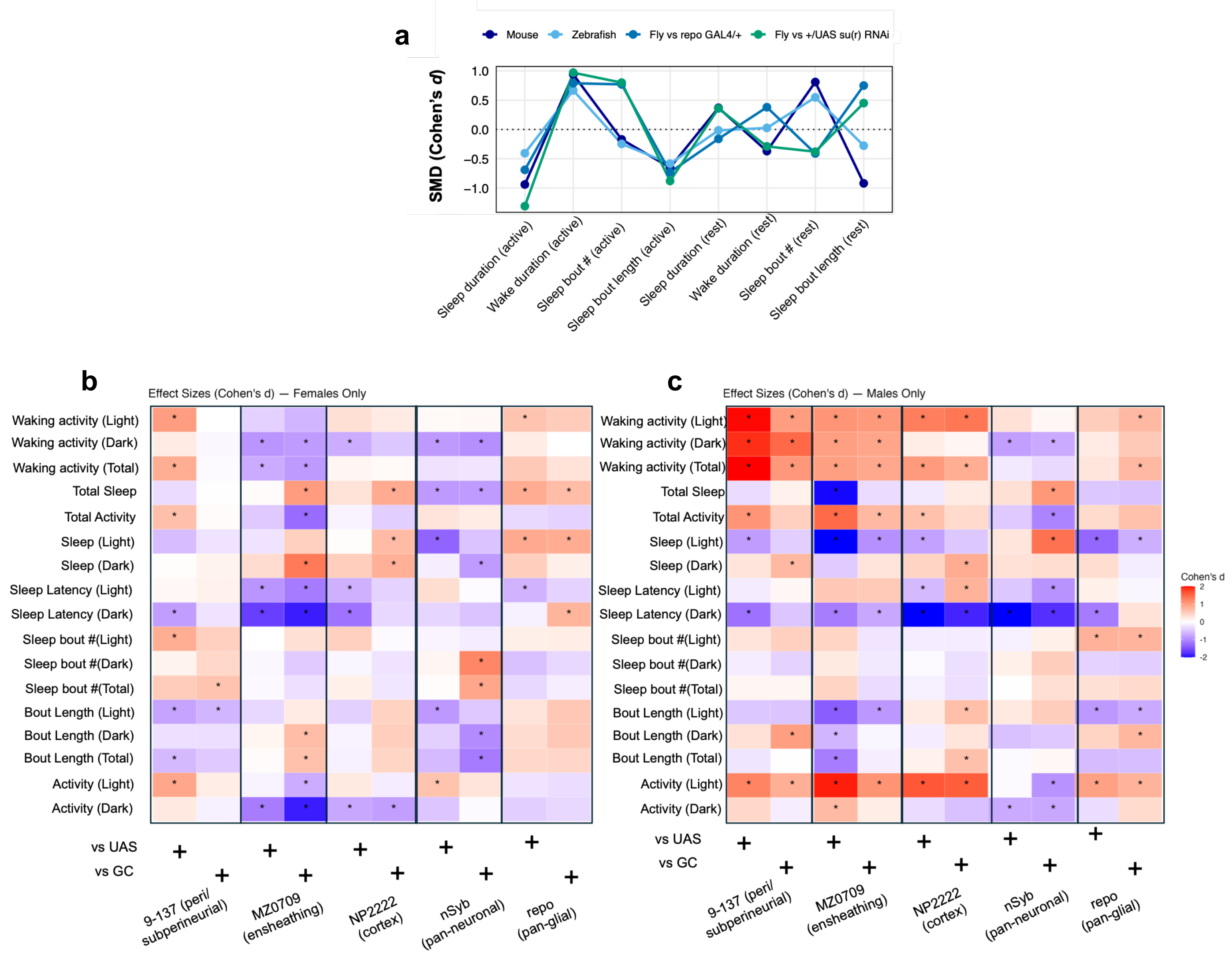


**Extended Data Fig. 11. Sleep and activity parameters following *Drosophila su(r)* knockdown. a.** Standardized mean difference (SMD; Cohen's *d*) comparison across mouse, zebrafish, and *Drosophila*. *Drosophila repo*-GAL4>UAS *su(r)* RNAi data are compared against both parental lines. All sleep and activity measures in females (**b**) and males (**c**). Data represent Cohen's *d* effect sizes relative to each parental control as indicated at the bottom. UAS = UAS RNAi-only control, GC = Gal4-only control. Asterisks indicate 95% confidence intervals not overlapping zero; however, no multiple comparisons corrections were applied.

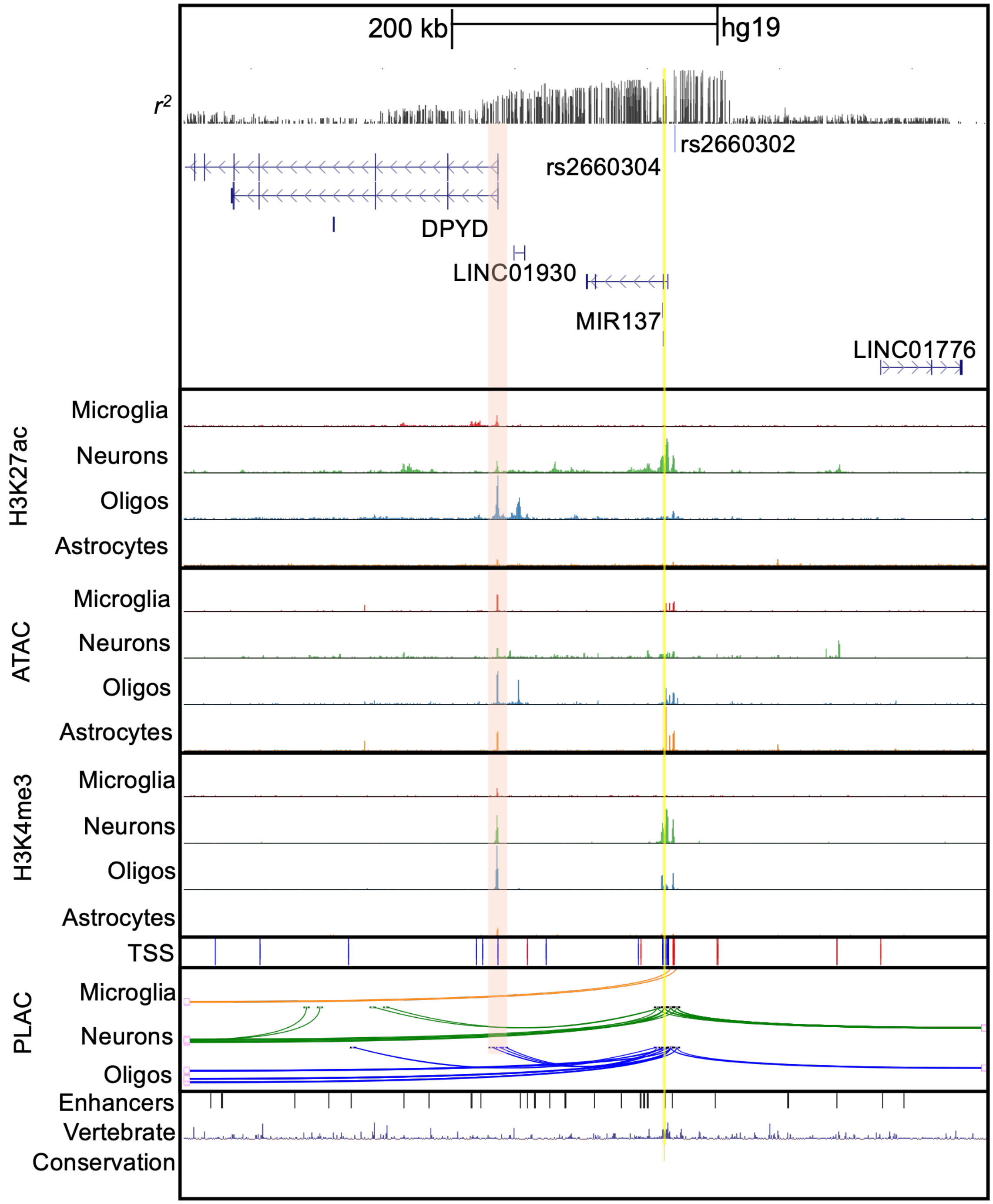


**Extended Data Fig. 12. Chromatin landscape at the *DPYD-MIR137HG* locus.** UCSC browser plot showing from the top: Linkage Disequilibrium ($r^2$ CEU from LDlink) for lead sleep duration GWAS variant rs2660302. Top ForgeDB scored regulatory variant, rs2660304, is indicated by the yellow line. Genes are labeled with arrows indicating strand direction. Orange highlight represents *DPYD* promoter region indicated by histone marks and transcription start site. H3K27ac, ATAC, H3K4me3, and PLAC-seq data from Nott et al. 2019 in different brain cell types. Oligos = oligodendrocytes. PLAC-seq = proximity ligation-assisted ChIP-seq showing chromatin interactions. All interactions are shown. TSS = transcription start sites. Fantom5 enhancers indicated by vertical black bars. Vertebrate conservation is shown. Note rs2660304 falls in a chromatin contact loop with the promoter of *DPYD* in oligodendrocytes. Each chromatin contact point shows accessible chromatin marks of promoters (H3K4me3 + ATAC + H3K27ac) and enhancers (H3K27ac + ATAC). Browser coordinates: (hg19) Chr1:98,149,243-98,756,598.
